# Altered cognitive processes shape tactile perception in autism

**DOI:** 10.1101/2025.06.25.661477

**Authors:** Ourania Semelidou, Mathilde Tortochot-Megne Fotso, Adinda Winderickx, Andreas Frick

## Abstract

Altered sensory perception is a hallmark of autism and shapes how individuals engage with their environment, with tactile perception playing a critical role in daily functioning and for social interactions. While sensory alterations are thought to contribute to cognitive differences in autism, the impact of cognition on sensory perception remains unclear. Here, we investigated how cognitive processes modulate tactile perception in the *Fmr1-*KO genetic mouse model of autism through a translational perceptual decision-making task. Our results revealed salience-dependent cognitive alterations that influenced sensory performance. During training, *Fmr1*^-/y^ male mice distinguishing between a high- and a low-salience stimulus exhibited an increased choice consistency bias in low-salience trials. When tested across a continuum of intermediate stimulus intensities, these mice demonstrated enhanced tactile discrimination of low-salience stimuli but reduced discrimination facilitation for stimuli crossing category boundaries. These effects were accompanied by diminished integration of sensory history and were dissociable from the attention deficits that emerged under high cognitive load. Together, our findings reveal that tactile perceptual alterations reflect context-dependent weighting and integration of sensory information during decision-making rather than uniform sensory deficits or enhancements, supporting a shift beyond traditional sensory–cognitive dichotomies.

## Introduction

Autism is a neurodevelopmental condition that affects approximately 1 in 31 children (Shaw et al., 2025). Autistic individuals display differences in social interaction and communication, repetitive behaviors, intense or special interests, and altered sensory experience as core symptoms (American Psychiatric Association, 2022). Sensory alterations strongly affect the way autistic individuals interact with their environment. According to the DSM-5-TR, these are broadly expressed as “*hyper- or hyporeactivity to sensory input or unusual interests in sensory aspects of the environment*” and are reported in 90% of autistic individuals, spanning all sensory modalities (Robertson and Baron-Cohen, 2017).

Amongst sensory modalities, touch is the first sense to develop and plays a fundamental role in the exploration of the environment, the definition of “self”, and early social bonding (Bremner and Spence, 2017). Tactile processing alterations have a significant impact on daily functioning and are thought to contribute to repetitive behaviors and social interaction difficulties in autism (Foss-Feig et al., 2012; Green et al., 2018; Thye et al., 2018; He et al., 2021a; Zhai et al., 2023; Carati et al., 2024). However, clinical studies focusing on tactile discrimination have yielded controversial results (Zetler et al., 2019), likely due to the substantial heterogeneity of tactile processing alterations in autism (Mikkelsen et al., 2018) and inconsistent terminology across studies (He et al., 2023). In contrast, numerous studies in the visual and auditory domains (Plaisted et al., 1998a; O’Riordan and Plaisted, 2001; O’Riordan et al., 2001; Bonnel et al., 2003; O’Riordan and Passetti, 2006; Samson et al., 2006; Rotschafer, 2021) have supported enhanced sensory processing in autism, as proposed by several influential models of the condition (Plaisted, 2001; Frith, 2003; Mottron et al., 2006).

Although clinical studies highlight the heterogeneity of tactile responses in autism, variability in experimental protocols and stimulus types has made it difficult to draw cohesive conclusions. Tactile discrimination of vibrotactile stimuli delivered to the glabrous skin of the fingers was reported reduced (Puts et al., 2014; Espenhahn et al., 2023), or intact (He et al., 2021a; Asaridou et al., 2022) in autistic individuals. Similarly, studies of sharp-dull discrimination have reported both reduced (Abu-Dahab et al., 2013) and typical performance (Minshew et al., 1997; Minshew and Hobson, 2008), as have studies of form discrimination (Minshew and Hobson, 2008; Demopoulos et al., 2015; Failla et al., 2017). Translational approaches employing mouse models of autism in combination with psychometric measures from well-controlled behavioral tasks may help clarify whether these inconsistencies reflect genuine inter-individual variability or arise from methodological differences.

While tactile discrimination has been explored in several clinical studies, it remains underexplored in mouse models of autism. Rodent studies leveraging innate responses have reported atypical whisker-dependent texture discrimination (Balasco et al., 2021, 2022; Mattioni et al., 2024) and forepaw-dependent roughness discrimination (Orefice et al., 2016). However, these studies have not employed psychophysical methods to quantify tactile discrimination, restricting the translatability of their findings. Implementing psychophysics in animal models might clarify previous inconsistencies between studies in mice (Orefice et al., 2016) and humans (O’Riordan and Passetti, 2006).

Sensory alterations are thought to impact cognition in autism (Haigh, 2018). However, sensory perception is shaped not only by the physical properties of stimuli but also by higher-order cognitive processes such as attention and categorization—both of which are altered in autism (Allen and Courchesne, 2001; Church et al., 2010; Gastgeb and Strauss, 2012; Cascio et al., 2015; Barbot et al., 2018; Rong et al., 2021a). Moreover, autistic individuals demonstrate differences in how prior beliefs and expectations (priors) are integrated with sensory input during perceptual decision-making. They tend to rely less on contextual priors (Amoruso et al., 2019) and more heavily on prior choices (Feigin et al., 2021), and often form priors that are imprecise or inflexible (Zaidel et al., 2015; Sapey-Triomphe et al., 2021b). Notably, variability in the use of priors has also been linked to individual differences in sensory responsivity (Sapey-Triomphe et al., 2021a). Thus, cognitive differences may compound or interact with primary sensory processing alterations, contributing to the distinct perceptual experiences observed in autism.

In this study, we investigated the interplay between tactile discrimination and cognitive processes during perceptual decision-making. To enhance translational relevance and address questions of inter-individual heterogeneity, we developed a forepaw-based 2-Alternative Choice task for mice, using vibrotactile stimuli analogous to those used in human psychophysical protocols. Leveraging this task, we explored how stimuli of varying salience are discriminated in the *Fmr1*^-/y^ mouse model of autism and disentangled stimulus-driven from cognitively modulated-tactile responses.

Our results revealed salience-dependent alterations in both tactile processing and cognitive modulation in *Fmr1*^-/y^ mice, with a decreased influence of cognitive processes on tactile perception. During training, *Fmr1*^-/y^ mice exhibited heightened choice consistency bias during low-salience trials, which contributed to reduced performance during perceptual learning. In contrast, trained *Fmr1*^-/y^ mice displayed enhanced discrimination of low-salience stimuli, but a diminished facilitatory effect of categorization on across-category discrimination. Despite their increased discrimination sensitivity at the low-salience range, *Fmr1*^-/y^ mice showed attention deficits specifically for these stimuli under high cognitive load conditions. While a strong choice consistency bias was observed during the testing phase for both WT and *Fmr1*^-/y^ mice, integration of recent sensory history was selectively disrupted in *Fmr1*^-/y^ mice, such that past sensory information no longer informed ongoing decisions.

These findings uncover distinct, salience-dependent cognitive alterations that shape tactile perception in the *Fmr1*^-/y^ model, advancing our understanding of tactile perception in autism. Importantly, our results indicate that the variability observed in clinical studies may arise not solely from primary sensory alterations but also from differences in cognitive context, attentional demands, or task complexity—offering a potential framework to reconcile prior inconsistencies in the field.

## Results

### *Fmr1^-^*^/y^ mice show intact learning rate and trajectory during perceptual learning

To investigate perceptual and cognitive alterations in a highly translational approach, we developed a forepaw-based 2-Alternative Choice perceptual decision-making task in the *Fmr1-*KO mouse model of autism. In this task, mice learn to discriminate between vibrotactile stimuli that differ in perceptual salience, which we define as the prominence of a stimulus based on its physical amplitude. To ensure that all mice could reliably detect the applied stimuli, we selected amplitudes above the perceptual thresholds previously established for both wild-type (WT) and *Fmr1*^-/y^ male mice (Semelidou et al., 2024). Accordingly, all stimuli used in the present study (40 Hz, ≥12 µm) were well above the detection threshold (10 Hz, 4.46 µm for WT; 10Hz, 7.29 µm for *Fmr1*^-/y^ mice), allowing us to specifically probe discrimination, categorization, and decision-making rather than sensory detection.

During each trial, head-fixed, water-controlled mice received a 500 ms vibrotactile stimulus to the forepaw at 40 Hz with either high amplitude (26 μm) or low amplitude (12 μm). Stimulus salience was defined in a relative manner: across animals, the lower-amplitude stimulus elicited consistently higher miss rates, indicating reduced perceptual strength and attentional engagement within the suprathreshold range. We therefore refer to the 12 μm stimulus as “low salience” and the 26 μm stimulus as “high salience.” Mice were trained to report high-salience stimuli by licking the right lick-port, and low-salience stimuli by licking the left port within a 2-s response window following stimulus delivery. Incorrect choices resulted in a 5-s timeout (Fig. 1A).

**Figure 1.**
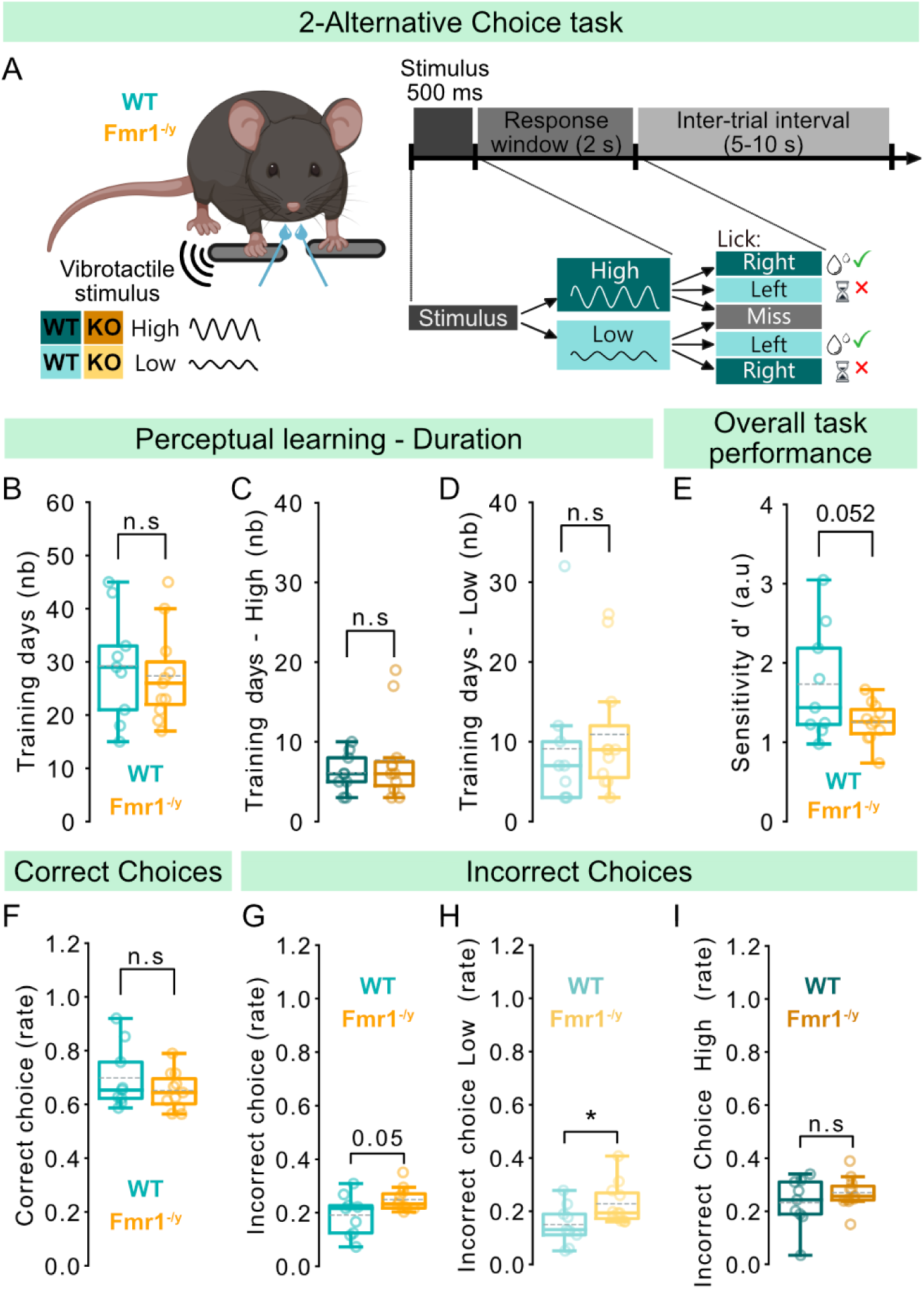
Perceptual learning performance in a forepaw-based decision-making task. For panels **B, C, D, E, F, G, H, I, J, K:** n=9 WT, 11 *Fmr1*^-/y^ male mice. **A,** Left: schema showing the behavioral setup. Right: Trail protocol and behavioral outcomes depending on the type of trial and the animal’s response. **B,** Total number of days spent in training for WT and *Fmr1*^-/y^ mice. **C,** Total number of days spent in training until the criterion was met for high-salience stimuli for WT and *Fmr1*^-/y^ mice. **D,** Total number of days spent in training until the criterion was met for low-salience stimuli for WT and *Fmr1*^-/y^ mice. **E,** Sensitivity d’ throughout the training period for WT and *Fmr1*^-/y^ mice. **F,** Correct choice rate for both high- and low-salience trails throughout the training period for WT and *Fmr1*^-/y^ mice. **G,** Incorrect choice rate for both high- and low-salience trails throughout the training period for WT and *Fmr1*^-/y^ mice. **H,** Incorrect choice rate for low-salience trials throughout the training period for WT and *Fmr1*^-/y^ mice. **I,** Incorrect choice rate for high-salience trails throughout the training period for WT and *Fmr1*^-/y^ mice. P values were computed using two-sided t-test for panels **B, E, F, G, I,**; Mann-Whitney test for panels **C, D, H**; *P < 0.05 or n.s, not significant.

During early training sessions, performance was low, characterized by a low proportion of correct and a high proportion of incorrect choices, which either remained stable or gradually improved over the course of the session (Fig. S1A-left). Licking patterns were also inconsistent during these early sessions and poorly time-locked to the response window (Fig. S1A-right). In contrast, late training sessions showed consistently high correct choice rates and low incorrect choice rates (Fig. S1B-left), along with stable licking patterns during the response window for both high- and low-salience trials (Fig. S1B-right).

Learning difficulties and altered learning trajectories have been reported in autistic individuals and replicated in several mouse models of autism, including *Fmr1*^-/y^ mice (Arnett et al., 2014a; Goel et al., 2018; Mercado et al., 2020; Khachadourian et al., 2023; Shenouda et al., 2023; Mol et al., 2024). To assess whether *Fmr1*^-/y^ mice exhibit alterations in learning rate in our task, we compared the number of days required to reach the predefined learning criterion (>70% correct choices for each stimulus salience across three consecutive training days). Our results showed no significant differences between *Fmr1*^-/y^ mice and their WT littermates in the total number of training days to reach criterion (Fig. 1B; T test: t = 0.438, p = 0.667) or in the duration spent at each stage of the training protocol (Fig. S1C-D, see Methods; Blocks: T test: t = 1.259, p = 0.224; Mixed: T test: t = -0.185, p = 0.855). Similarly, both genotypes required a comparable number of days to learn the association between stimulus salience and the correct lick-port (Fig. 1C-D; High salience: Mann-Whitney U test: U=45.5, p = 0.788; Low salience: Mann-Whitney U test: U = 38.5, p = 0.422).

To capture potential genotype differences in the learning dynamics during training, we analyzed the performance of all mice that reached the learning criterion during the training phase with mixed low- and high-salience trials. Both WT and *Fmr1*^-/y^ mice showed similar learning dynamics, with a slow, gradual improvement across days (main effect of day: β = 0.007, SE = 0.004, z = 1.882, p = 0.060, 95% CI [0.00, 0.01]) (Fig. S2A-top). Consistent with the extended training duration required for the task, learning slopes were shallow and comparable between genotypes (Fig. S2B; Mann-Whitney U test: U=38.0, p=0.536). Importantly, the Genotype × Day interaction was small (β = 0.002), indicating no detectable genotype differences in learning rate across animals.

These similar learning trajectories were accompanied by comparable baseline performance on the first day of training across genotypes (Fig. S2A-top, S2C; genotype effect: β = −0.087, SE = 0.066, z = -1.328, p = 0.184, p = 0.184, 95% CI [-0.22, 0.04]) and similar within-animal day-to-day performance variability (Fig. S2A-bottom; Fig. S2D; T test: t=-0.302, p=0.767).

We next assessed whether performance differed during an intermediate stage of learning, defined for each animal as the middle three days of its training period. Here, both genotypes again showed comparable performance (Fig. S2E, slope; T-test: t = 0.801, p = 0.437) with similar correct-choice rates (Fig. S2F; T-test: t=1.240, p=0.228), indicating no genotype-specific differences during intermediate learning.

Together, these results show that *Fmr1*^-/y^ mice display an intact learning rate and trajectories in this 2-AFC perceptual decision-making task.

### *Fmr1^-^*^/y^ mice display impaired performance on low-salience trials during perceptual learning

Given that comparable learning trajectories and training duration to reach the criterion do not exclude the presence of performance alterations, we next evaluated additional behavioral metrics during training with mixed high- and low-salience trials. Specifically, we analyzed overall discrimination sensitivity (d’) as well as correct and incorrect choice rates throughout training to determine whether *Fmr1*^-/y^ mice exhibited altered task performance despite intact learning progression.

To quantify mice’s sensitivity in discriminating high-versus low-salience stimuli throughout training, we used the signal detection theory and computed the sensitivity index d′. This measure captures stimulus discriminability by comparing normalized hit rates (rightward licks on high salience trials) and false alarm rates (rightward licks on low salience trials). *Fmr1*^-/y^ mice showed a strong trend with a large effect size toward reduced overall task sensitivity compared to WT littermates (Fig. 1E; T-test: t=-2.083, p=0.052, Hedges’ g=0.90), indicating diminished perceptual performance during training. These results demonstrate that, despite acquiring the task at a comparable rate (Fig. 1B-D) and with similar learning trajectories (Fig. S2), *Fmr1*^-/y^ mice display a trend for diminished performance during salience discrimination training.

To complement our sensitivity (d′) analysis, we also quantified correct and incorrect choice rates, allowing us to assess both rightward and leftward lick responses. Correct choice rates were comparable between *Fmr1*^-/y^ and WT mice (Fig. 1F, T-test: t=1.064, p=0.307) with similar Hit rates for both high- and low-salience stimuli (Fig. S1E–F, High salience-T-test: t=0.721, p=0.480; Low salience-T-test: t=1.060, p=0.303). In contrast, *Fmr1*^-/y^ mice exhibited a significantly higher rate of incorrect choices (Fig. 1G; T-test: t=-2.099, p=0.050; Hedge’s g=0.90), an effect that was specific to low-salience trials (Fig. 1H; Mann–Whitney U = 23.0, p = 0.048; Hedge’s g = 0.97), while high-salience error rates were similar across genotypes (Fig. 1I; T-test: t=-1.072; p=0.298). After outlier analysis and exclusion of one *Fmr1*^-/y^ mouse, the genotype difference was reduced to a trend, though with a large effect size, for incorrect choices (T-test: t=-1.803; p=0.089, Hedge’s g = 0.79) and for incorrect choices during low-salience trials (Mann–Whitney U = 23.0, p = 0.079; Hedge’s g = 0.89).

These results suggest a modest reduction in task performance in *Fmr1*^-/y^ mice, primarily driven by increased errors on low-salience trials.

### *Fmr1^-^*^/y^ mice show higher choice consistency bias during training

Perceptual learning can further reveal potential alterations in decision strategy, manifested as changes in response bias (decision criterion c) or expectation bias (priors). To examine whether mice exhibited stereotyped licking toward one of the two lickports—a potential indicator of response bias—we calculated the decision criterion. This analysis revealed no significant differences between *Fmr1*^-/y^ mice and WT littermates (Fig. 2A; Mann–Whitney U: U=65.000, p=0.254), with both groups showing a bias toward the left lickport (WT: Wilcoxon signed-rank test W=5.000, p=0.039; *Fmr1*^-/y:^ Wilcoxon signed-rank test W=10.000, p=0.042).

**Figure 2.**
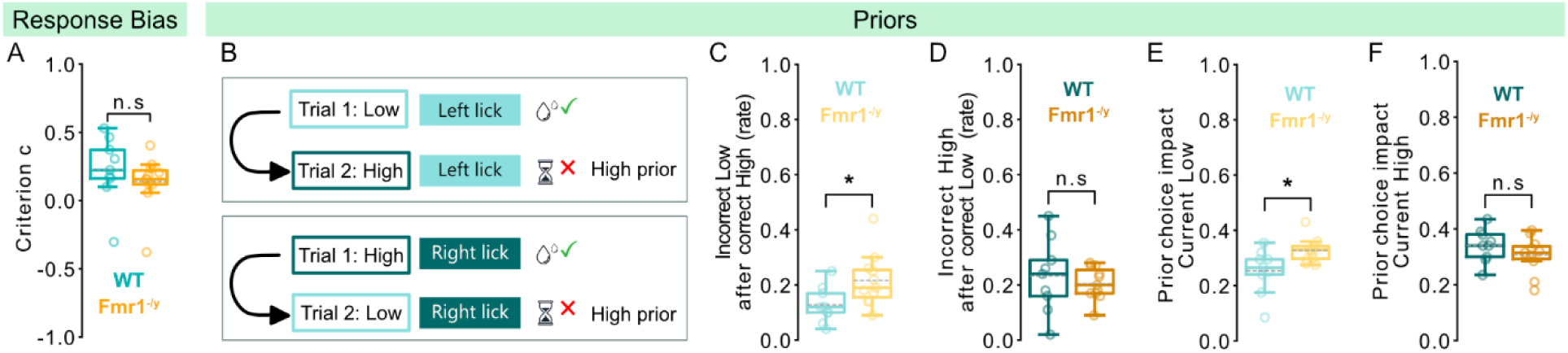
Overall strategy and impact of prior choice during perceptual learning. For panels **A, C, D, E, F:** n=9 WT, 11 *Fmr1*^-/y^ male mice. **A,** Criterion depicting the licking strategy of the animals. **B,** Schema showing an example of how high impact of prior choice on the current trial affects the response during a high-salience trial (top) or low-salience trial (bottom). **C,** Proportion of incorrect responses in low-salience trials immediately following a correctly rewarded high-salience trial. **D,** Proportion of incorrect responses in high-salience trials immediately following a correctly rewarded low-salience trial. **E,** Proportion of correct responses in low-salience trials immediately following a correctly rewarded low-salience trial and incorrect responses in low-salience trials immediately following a correctly rewarded high-salience trial. Rates are corrected over the total number of correct and incorrect choices in low-salience trials. **F,** Proportion of correct responses in high-salience trials immediately following a correctly rewarded high-salience trial and incorrect responses in high-salience trials immediately following a correctly rewarded low-salience trial. Rates are corrected over the total number of correct and incorrect choices in high-salience trials. P values were computed using Mann-Whitney test for panel **A,;** two-sided t-test for panels **C, D, E, F,**; *P < 0.05 or n.s, not significant.

Since mice did not exhibit differences in stereotyped licking, we could rule out simple response biases and further assess whether their decisions are influenced not only by the sensory features of stimuli but also by previous experiences that shape expectations (i.e., priors). To investigate the impact of prior experience on current choices, we analyzed behavioral responses on a trial-by-trial basis. We first examined how a rewarded choice in the previous trial influenced the performance in the current trial (Fig. 2B). *Fmr1*^-/y^ mice exhibited an increased rate of incorrect responses to low-salience stimuli when these were preceded by a rewarded high-salience stimulus (Fig. 2C; T-test: t=-2.352, p=0.030). In contrast, no genotype difference was observed when a high-salience stimulus followed a rewarded low-salience trial (Fig. 2D; T-test: t=0.603, p=0.554).

We next tested whether *Fmr1*^-/y^ mice were more influenced by their prior choices. To assess this, we compared the rates of repeated prior choices in current low- and high-salience trials, controlling for the total number of correct and incorrect responses. Our analysis revealed higher previous choice repetition during low-salience trials in *Fmr1*^-/y^ compared to WT mice (Fig. 2E; T-test: t=-2.618, p=0.017), but a comparable impact of prior choices during high-salience trials (Fig. 2F; T-test: t=1.215, p=0.240).

To verify that the observed differences in performance between WT and *Fmr1*^-/y^ mice reflected differential reliance on priors rather than differences in prior strength per se, we quantified the strength of the priors induced by high- and low-salience stimuli. Specifically, we calculated the rate of choices that were repeated following a low- or high-salience trial, correcting for the overall rate of correct and incorrect responses. Our results showed similar prior strength between groups, both for high-salience (Fig. S3A; T-test: t=-1.705, p=0.105) and for low-salience stimuli (Fig. S3B; Mann–Whitney U: U=38.500, p=0.425).

Together, these results indicate an increased choice consistency bias during low-salience trials, which contributes to performance differences during perceptual learning in *Fmr1*^-/y^ mice.

### *Fmr1^-^*^/y^ mice do not show attention deficits during perceptual learning

Stimulus salience is closely linked to attentional processes (Kerzel and Schönhammer, 2013; Forschack et al., 2022; Bouvier et al., 2023), and attentional alterations are well-documented in autistic individuals (Allen and Courchesne, 2001; Grubb et al., 2013; Barbot et al., 2018; Licznerski et al., 2020; Rong et al., 2021b). Given that *Fmr1*^-/y^ mice exhibit salience-dependent performance differences during training (Fig. 1-2), we asked whether their reduced task performance might also reflect underlying attention deficits. To assess attention, we analyzed the trials in which the animal failed to respond to the stimulus (Miss trials) (Fig 1A). Within-genotype analysis confirmed studies linking salience and attention in humans (Kerzel and Schönhammer, 2013), and validated that low amplitude stimuli are less salient, yielding significantly higher Miss rates in both groups (Fig. S3C; WT: Wilcoxon signed-rank: W=0.000, p=0.004; *Fmr1*^-/y^: Paired T-test: t=-4.113, p=0.002). However, there were no genotype differences in Miss rates for either high- or low-salience stimuli (Fig. S3D-E; Miss rate High: Mann–Whitney U: U=51.000, p=0.939; Miss rate Low: Mann–Whitney U: U=50.000, p=1.000). These data demonstrate that low-salience stimuli are less effective at capturing attention in both *Fmr1*^-/y^ and WT mice, and that the reduced task performance of *Fmr1*^-/y^ mice during perceptual training cannot be attributed to an attentional deficit during low-salience trials.

### Trained *Fmr1^-^*^/y^ mice show enhanced discrimination of low-salience stimuli

Altered tactile discrimination has been reported in clinical studies, though findings are inconsistent (reviewed in (Zetler et al., 2019)), and has also been observed in mouse models of autism (Orefice et al., 2016; Balasco et al., 2021, 2022; Mattioni et al., 2024). To assess tactile discrimination in our study, we incorporated the original training stimuli along with 6 intermediate amplitudes, spaced 2 µm apart. Stimuli ranging from 20 to 26 µm (20, 22, 24, 26 µm) were categorized and rewarded as high-salience stimuli, while those from 12 to 18 µm (12, 14, 16, 18 µm) were treated as low-salience stimuli (Fig. 3A). Only animals that successfully acquired the task during the training phase were included in the tactile discrimination test. *Fmr1*^-/y^ and WT mice showed similar performance during the last three days of training, with no significant differences in task sensitivity (d’; T-test: t=0.800, p=0.438; Fig. S4A), rates of correct (Fig. S4B-C; High: T-test: t=0.442, p=0.666; Low: T-test: t=0.697, p=0.498) and incorrect responses (Fig. S4D-E; High: T-test: t=-0.680, p=0.509; Low: T-test: t=-1.069, p=0.305), decision bias (Fig. S3F; criterion c; Mann–Whitney U: U=28.000, p=0.955), use of priors (Fig S4G-L; Fig. S4G: T-test: t=0.485, p=0.636; S4H: T-test: t=0.155, p=0.879; S4I: T-test: t=-0.667, p=0.516; S4J: T-test: t=-0.667, p=0.516; S4K: T-test: t=-0.072, p=0.944; S4L: T-test: t=-0.472, p=0.644), or attention (Fig. S4M-N; High: Mann–Whitney U: U=32.500, p=0.555; Low: T-test: t=0.087, p=0.932).

**Figure 3.**
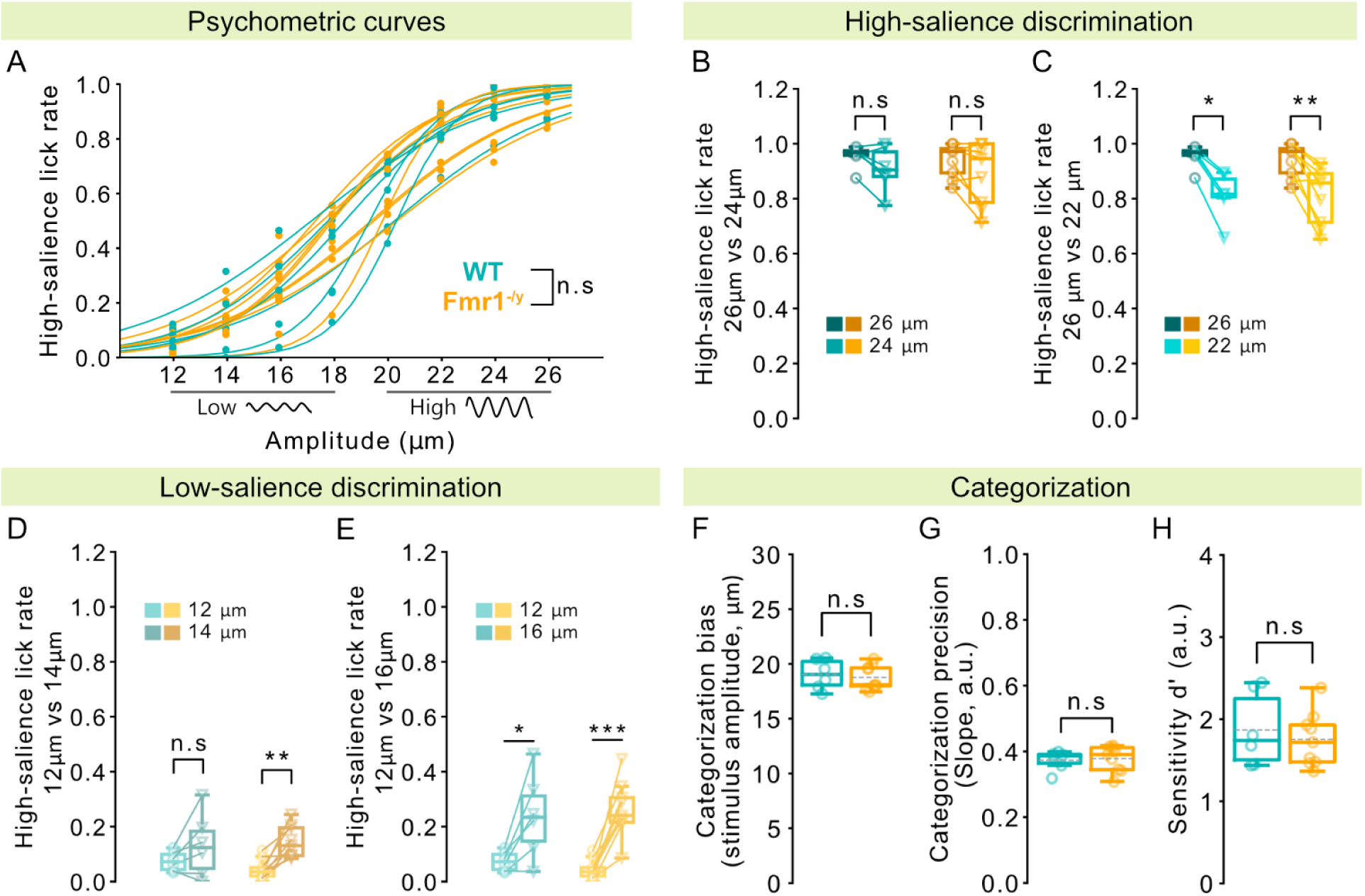
Tactile discrimination and categorization. n=6 WT, 9 *Fmr1*^-/y^ male mice. **A,** Psychometric curves for WT and *Fmr1*^-/y^ mice generated based on the high-salience lick rate (rate of rightward licks) across 8 different amplitudes. Stimuli between 12–18 µm were designated as low-salience and rewarded at the left lickport, while stimuli between 20–26 µm were designated as high-salience and rewarded at the right lickport. Each amplitude was presented an average of 84 times. **B,** Comparison of high-salience (rightwards) responses for high-salience stimuli of 26 and 26 µm. **C,** Comparison of high-salience (rightwards) responses for high-salience stimuli of 26 and 22 µm. **D,** Comparison of high-salience (rightwards) responses for low-salience stimuli of 12 and 14 µm. **E,** Comparison of low-salience (leftwards) responses for low-salience stimuli of 12 and 14 µm. **F,** Categorization bias calculated based on the psychometric curves. **G**, Categorization precision computed based on the slope of the psychometric curves. **H,** Sensitivity d’ of the responses in all stimulus amplitudes. P values were computed using Mixed Linear Model Regression for panel **A,**; two-sided paired t-test for panels **B-right, C-right, D, E,;** two-sided t-test for panels **F, G, H,**; Wilcoxon signed-rank test for panels **B-left, C-left,;** Bonferroni correction was applied for panels **B, C, D,** **P < 0.01, *P < 0.05, or n.s, not significant.

To evaluate task performance, we quantified the rate of right-lick responses, corresponding to reports of high-salience stimuli, and constructed psychometric curves, which increased with stimulus amplitude in both genotypes. Mixed-effects modeling revealed a strong main effect of amplitude (all amplitude coefficients p < 0.005) but no evidence for a genotype-dependent change across the psychometric function (amplitude x genotype interaction coefficients p > 0.266), indicating comparable performance between WT and KO animals across stimulus intensities (Fig. 3A). A genotype-by-amplitude interaction was observed only at 18 µm amplitude (β = 0.127, SE = 0.06, z = -2.18, p = 0.030, 95% CI [-0.24, -0.01]), where responses were slightly reduced in WT mice. However, this effect did not survive post-hoc analysis (T-test: t = 0.568, p = 0.580) and did not generalize across amplitudes.

We then assessed the animals’ ability to discriminate high-salience stimuli by comparing the proportion of right-lick responses to 26 µm and 24 µm stimuli. Our results showed that neither WT nor *Fmr1*^-/y^ mice were able to reliably distinguish between these two amplitudes (Fig. 3B; WT: Wilcoxon signed-rank: statistic=3.000, p=0.156; *Fmr1*^-/y^: Paired t-test: t = 1.447, p = 0.221). However, both groups reliably discriminated between 26 µm and 22 µm stimuli (Fig 3C; WT: Wilcoxon signed-rank: statistic = 0.000, p = 0.031; *Fmr1*^-/y^: Paired t-test: t = -4.929, p = 0.001). These results demonstrate that high-salience stimuli differing by 4 µm in amplitude can be discriminated by both genotypes.

We next investigated low-salience stimulus discrimination by comparing the rate of right-lick responses (reporting high salience) to stimuli of 12 µm and 14 µm amplitude. WT mice responded similarly to both amplitudes (Fig. 3D-left; Paired t-test: statistic = -1.684, p = 0.153), consistent with their performance on high-salience stimuli differing by 2 µm (Fig. 3B-left). In contrast, *Fmr1*^-/y^ mice exhibited enhanced discrimination for these low-salience stimuli, with increased high-salience report rates for the 14 µm stimuli compared to those for 12 µm (Fig. 3D-right; Paired t-test: t = -5.007, p = 0.001). Both genotypes efficiently discriminated 12 µm from 16 µm stimuli (Fig. 3E; WT: Paired t-test: t = -3.274, p = 0.022; *Fmr1*^-/y^: Paired t-test: t = -6.772, p < 0.001). This enhanced discrimination in *Fmr1*^-/y^ mice was further supported by low-salience reports that showed lower low-salience lick rates for 14 µm than 12 µm stimuli (Fig. S5A-right; Paired t-test: t = 2.394, p = 0.044), whereas WT mice responded similarly (Fig. S5A-left; Paired t-test: t = 1.589, p = 0.173), validating that improved discrimination was not driven by higher miss rates for 12 µm stimuli.

Together, these results demonstrate that *Fmr1*^-/y^ mice present enhanced tactile fine-discrimination for low-salience stimuli.

### *Fmr1^-^*^/y^ mice exhibit intact salience categorization

In addition to assessing stimulus discrimination, our task allowed us to evaluate whether *Fmr1*^-/y^ and WT mice form the same categories of low- and high-salience stimuli. Using the psychometric curves of the animals’ high-salience responses (Fig. 3A), we measured the categorization bias and precision. No differences were observed between *Fmr1*^-/y^ mice and their WT littermates regarding their categorization bias (Fig. 3F; T-test: t=0.432, p=0.673) or precision (slope, Fig. 3G; T-test: t=-0.283, p=0.782), showing comparable salience categorization between groups. These findings were further confirmed by similar overall task sensitivity (d’, Fig. 3H; T-test: t=0.571, p=0.578). Furthermore, *Fmr1*^-/y^ mice showed similar rates of correct (Fig. S5B-C; High: T-test: t=-0.160, p=0.875; Low: T-test: t=1.389, p=0.188) and incorrect choices (Fig. S5D-E; High: T-test: t=0.421, p=0.681; Low: T-test: t=-0.709, p=0.491), as well as comparable overall decision strategy (Fig. S5F, criterion c; T-test: t=0.639, p=0.533) compared to their WT littermates. In conclusion, these results demonstrate similar categorization of high- and low-salience stimuli in *Fmr1*^-/y^ and WT mice.

### *Fmr1^-^*^/y^ mice show reduced impact of categorization on stimulus discrimination

Higher-order processes such as categorization can amplify perceptual differences between stimuli belonging to different categories (Goldstone, 1994). Alterations in this behavioral process have been reported in autistic individuals, who, despite exhibiting intact visual categorization (in line with our findings in the tactile domain, Fig. 3F-H), show a reduced influence of categorization on perceptual discrimination that leads to diminished facilitation for stimuli belonging to different categories (Soulières et al., 2007). To examine whether salience categorization differentially impacts tactile discrimination in WT and *Fmr1*^-/y^ mice, we compared the discrimination accuracy for stimulus pairs that differ by 2 µm and either span across different salience categories (across-category, i.e., 18 µm and 20 µm stimuli) or fall within the same category. Discrimination accuracy was calculated as the difference in rightward choices between each pair of stimuli.

In WT mice, discrimination accuracy was higher for across-category stimulus pairs than for both within-category low- (Fig. 4A-left; Paired t-test, t = 4.452, p=0.007) and high-salience pairs (Fig. 4B-left; Paired t-test, t = 5.453, p = 0.003), indicating a robust facilitatory effect of categorization. *Fmr1*^-/y^ mice, in contrast, did not show this facilitation for low-salience stimuli, exhibiting a trend with medium effect size toward improved discrimination for across-category versus within-category low-salience pairs (Fig. 4A-right; Wilcoxon signed-rank: stat = 6.000, p = 0.055; Paired Hedges’ g = 0.640). For high-salience stimuli, however, *Fmr1*^-/y^ mice performed similarly to WT, showing enhanced discrimination for across-category pairs relative to within-category high-salience comparisons (Fig. 4B-right; Paired t-test, t = 3.496, p = 0.008).

**Figure 4.**
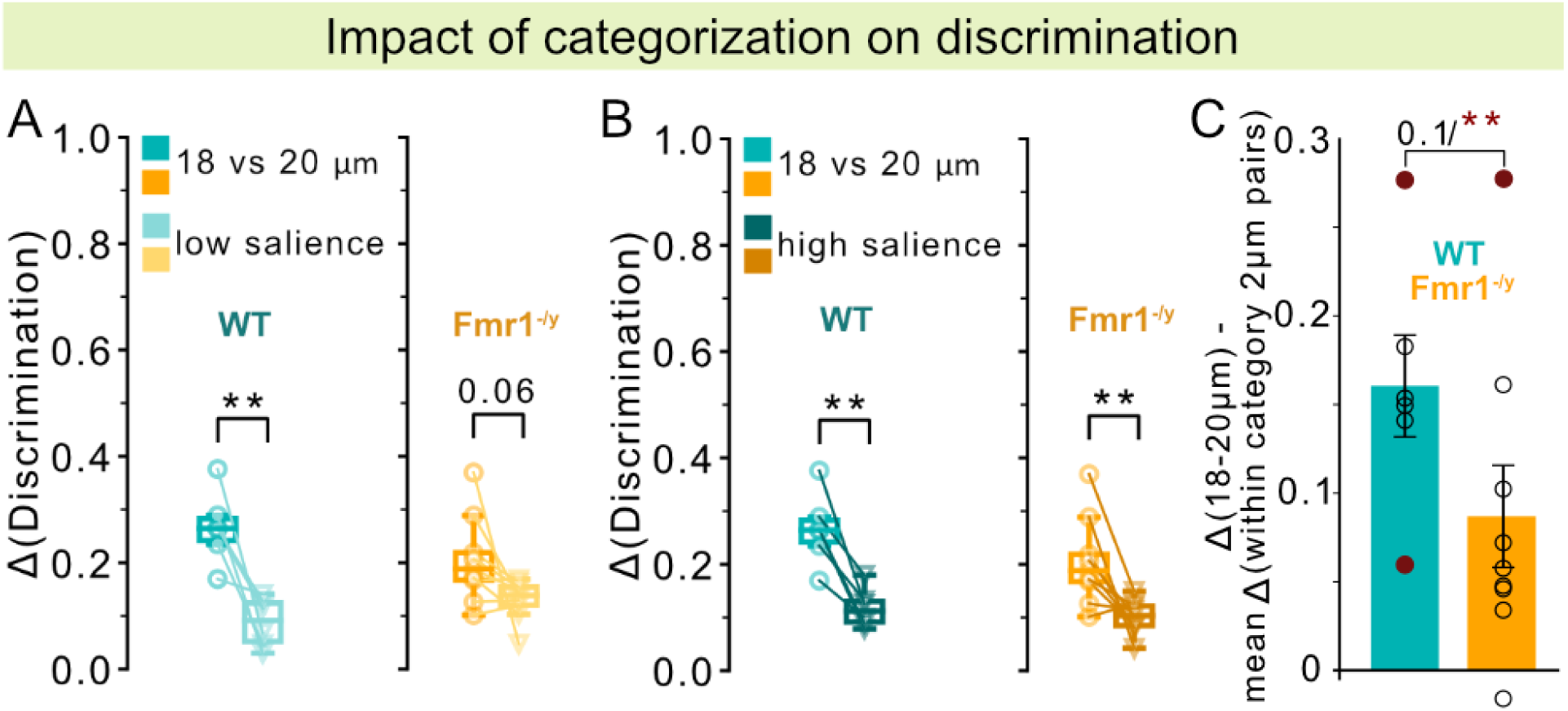
Impact of categorization on across-categories and within-category discrimination. n=6 WT, 9 *Fmr1*^-/y^ male mice. Delta discrimination accuracy, calculated as the difference in the rate of high-salience licks across stimulus pairs. **(A)** Delta discrimination accuracy between across-category stimuli (18 µm vs 20 µm) and within-category low-salience stimulus pairs, averaged as “low salience.” **B,** Same as (**A**,) but for within-category high-salience stimulus pairs. **C,** Delta discrimination accuracy across all stimulus pairs within the low-salience category. **D,** Same as (**C**,) but across all stimulus pairs within the high-salience category. P values were computed using two-sided paired t-tests for panels **A-left**, **B;** Wilcoxon signed-rank test for panel **A-right**, and two-sided t-test for panel **C,;** **P < 0.01, or n.s, not significant.

To directly compare WT and *Fmr1*^-/y^ mice, we quantified the categorical facilitation effect as the difference in discrimination accuracy between across- and within-category (high and low-salience) stimulus pairs. *Fmr1*^-/y^ mice showed a trend with a strong effect size toward reduced facilitation (T test: t = 1.81, p = 0.095; Hedge’s g = 0.86), which became significant after exclusion of outliers (Fig. 4C; T test: t = 4.57, p = 0.001; Hedge’s g = 1.94).

Together, these findings suggest that reduced categorization influence in *Fmr1*^-/y^ mice diminishes discrimination facilitation for stimuli belonging to different categories, revealing a genotype-specific alteration in how categorical information modulates tactile perception.

### High cognitive load conditions reveal salience-dependent attention deficits in *Fmr1*^-/y^ mice

Our results during perceptual learning revealed comparable, salience-dependent attention in WT and *Fmr1*^-/y^ mice (Fig. S3D-E). During the tactile discrimination/categorization phase of our task, animals were exposed to a high cognitive load, as they needed to discriminate and categorize eight stimulus intensities to obtain a water reward. To assess whether *Fmr1*^-/y^ mice exhibit attentional deficits in these high cognitive load conditions, we analyzed the rate of Miss trials as a proxy for attention.

While miss rates for high-salience stimuli did not differ between genotypes (Fig. 5A; all genotype x amplitude interactions p > 0.296), *Fmr1*^-/y^ mice showed elevated miss rates for low-salience stimuli during categorization (Fig. 5B). A mixed-effects linear model revealed that *Fmr1*^-/y^ mice had significantly higher miss rates than WT controls at the reference amplitude of 12 µm (β = 0.067, SE = 0.024, z = 2.77, p = 0.006, 95% CI [0.019, 0.114]). The genotype differences at 14 µm (β = −0.015, SE = 0.021, z = −0.70, p = 0.483) and 16 µm (β = −0.029, SE = 0.021, z = −1.37, p = 0.171) did not differ significantly from the effect observed at 12 µm. However, a significant genotype × amplitude interaction was observed at the highest amplitude (18 µm; β = −0.045, SE = 0.021, z = −2.16, p = 0.031, 95% CI [−0.087, −0.004]), indicating that genotype differences diminished the highest stimulus intensity of the low-salience range. Post-hoc comparisons further indicated a trend for increased misses at 12 µm with a large effect size (T-test: t = -2.8437, p = 0.058, Hedge’s g = 1.23), but showed no significant differences for the other low-salience stimuli (p > 0.21). Notably, during the training period—when cognitive load was lower—miss rates were comparable across genotypes (Fig. S3D-E).

**Figure 5.**
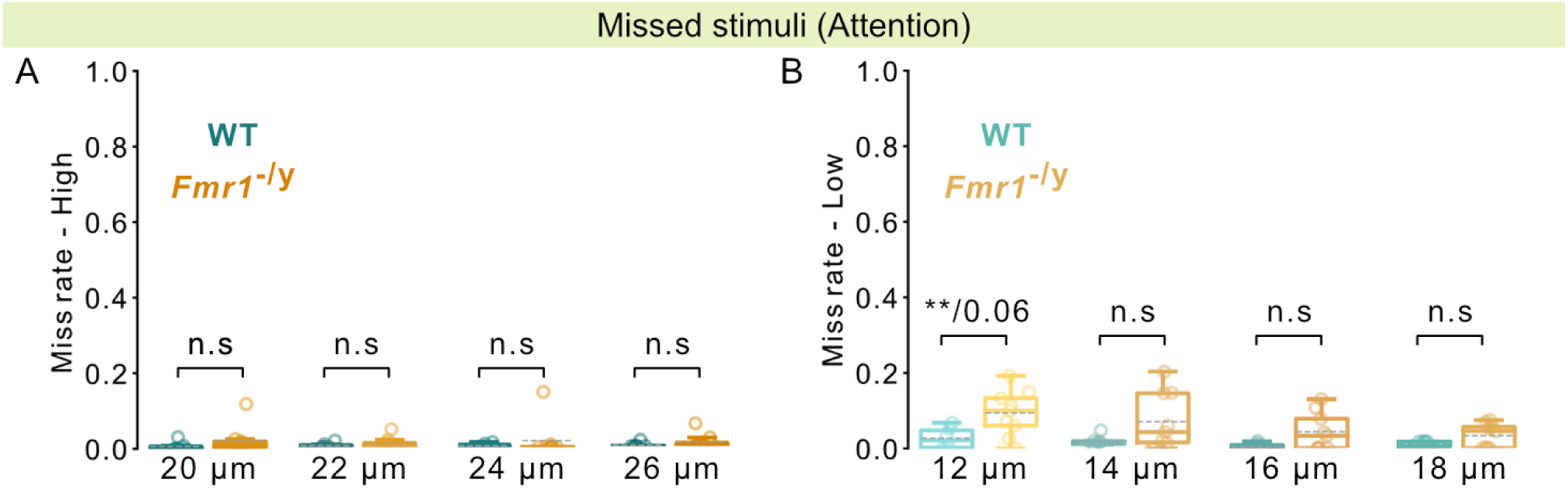
Attention in perceptual decision-making under high cognitive load. n=6 WT, 9 *Fmr1*^-/y^ male mice. **A,** Proportion of missed trials for high-salience stimuli. **B**, Proportion of missed trials for low-salience stimuli. P-values were obtained using mixed-effects linear models accounting for repeated measures across stimulus amplitudes and animals for panels **A, B**,; Post-hoc two-sided t-test for panel **B,-**12µm (p = 0.06); **P < 0.01, or n.s, not significant.

Together, these results reveal a salience-specific attentional deficit in *Fmr1*^-/y^ mice under high cognitive load, with low-salience stimuli being particularly vulnerable.

### *Fmr1*^-/y^ mice show preserved choice perseveration but disrupted sensory history integration during tactile categorization

We next examined how trial history—including prior stimulus, choice, and outcome—shapes current decisions during tactile categorization. To do so, we used generalized linear models (GLMs) with a binomial link function to predict high-salience licks (right-lick choices) based on the current stimulus, trial history, genotype, and their interactions. We first fit a main-effects model including the current stimulus, previous stimulus, previous outcome, previous choice, and genotype, and then extended this model to assess genotype-specific modulation of history effects through interactions between genotype and previous choice, outcome, and stimulus (Fig. 6a).

**Figure 6.**
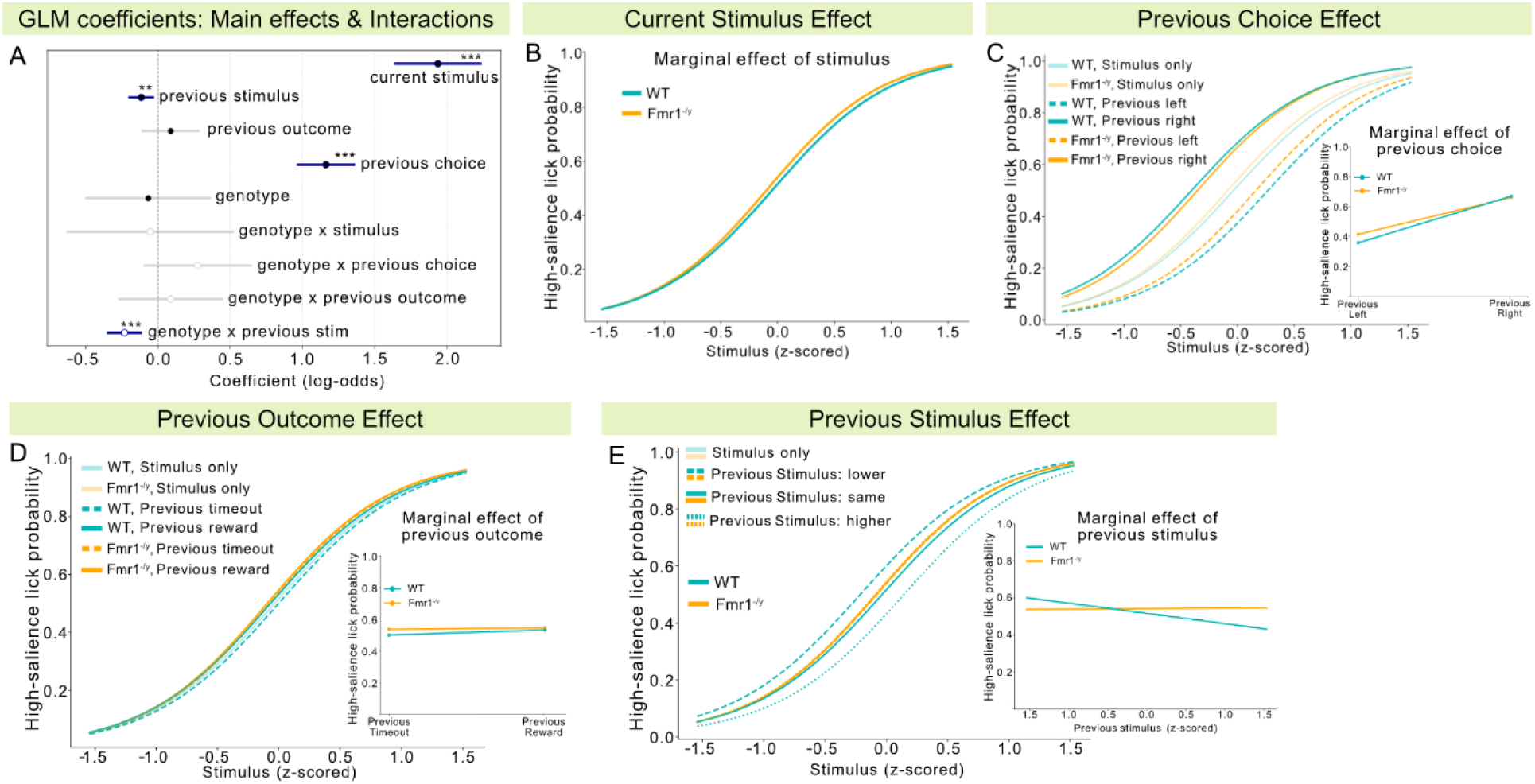
Trial history integration during tactile categorization. n=6 WT, 9 *Fmr1*^-/y^ male mice. **(A)** Generalized linear model (GLM) coefficients from binomial regression predicting high-salience (right-lick) choices based on current stimulus amplitude, previous stimulus amplitude, previous outcome, previous choice, genotype, and their interactions. Filled circles indicate coefficients from the main-effects model; open circles indicate coefficients from the extended model including genotype × history interactions. Error bars represent ±95% confidence intervals. Dark blue lines denote statistically significant effects. **(B)** Psychometric functions showing the effect of the z-scored current stimulus amplitude for WT and *Fmr1*^-/y^ mice. **(C)** Psychometric curves conditioned on the previous choice (left vs. right lick) for WT and *Fmr1*^-/y^ mice. **(D)** Psychometric curves conditioned on previous trial outcome (reward vs. timeout). **(E)** Psychometric curves conditioned on the intensity of the previous stimulus (z-scored). P values were computed using Mixed Linear Model Regression. ***P < 0.001, **P < 0.01.

As expected, the current stimulus was the dominant driver of choice in both genotypes (Fig. 6A-B; β = 1.94, SE = 0.15, z = 12.96, p < 0.001, 95% CI [1.64, 2.23]). However, previous choice also strongly influenced behavior (Fig. 6A). Both genotypes exhibited perseveration, as psychometric curves conditioned on the previous choice were systematically shifted following both prior right and left choices, indicating a robust tendency to repeat prior responses (Fig. 6A-C; β = 1.16, SE = 0.10, z = 11.85, p < 0.001, 95% CI [0.97, 1.36]). In contrast to the increased choice consistency bias observed during training (Fig. 2C-E), WT and *Fmr1*^-/y^ mice exhibited comparable levels of perseveration during categorization (genotype x previous choice interaction; β = 0.27, SE = 0.19, z = 1.47, p = 0.142, 95% CI [-0.09, 0.64]).

In contrast, previous outcome (reward or timeout) had a negligible effect on current choice in either genotype (Fig. 6A, D; β = 0.09, SE = 0.10, z = 0.89, p = 0.373, 95% CI [-0.11, 0.28]). Interestingly, the amplitude of the previous stimulus influenced decision-making (Fig. 6A; β = -0.12, SE = 0.04, z = -2.82, p = 0.005, 95% CI [-0.20, -0.04]) in a genotype-specific manner (Fig. 6A, E; genotype x previous stimulus interaction, β = -0.23, SE = 0.06, z = -4.03, p < 0.001, 95% CI [-0.34, -0.12]). In WT mice, larger-amplitude stimuli on the preceding trial slightly reduced the probability of choosing right on the subsequent trial, whereas lower-amplitude stimuli increased it (Fig. 6E). In contrast, choices of *Fmr1*^-/y^ mice were not modulated by the amplitude of the previous stimulus.

Together, these results show that tactile decisions in both genotypes are dominated by current sensory evidence and strongly biased by prior choices. While choice perseveration is preserved in *Fmr1*^-/y^ mice, integration of recent sensory history is selectively disrupted, revealing a genotype-specific deficit in how past sensory information informs ongoing decision-making.

## Discussion

Here, we developed a decision-making task based on psychophysics to dissociate stimulus-driven from cognitively-modulated tactile responses in the *Fmr1*^-/y^ mouse model of autism. Our findings reveal salience-dependent cognitive alterations that shape sensory performance. During perceptual learning, *Fmr1*^-/y^ mice exhibited an increased choice consistency bias during low-salience trials, which contributed to reduced task performance. During discrimination, *Fmr1*^-/y^ mice displayed an enhanced tactile sensitivity under low-salience conditions alongside decreased facilitation of across-category discrimination. Despite this increased sensitivity, these mice showed low salience-specific attentional impairments under high cognitive load, as well as faster updating of their world model with a decreased impact of sensory history on their current choice. Our findings highlight the interplay between sensory and cognitive alterations in autism, emphasizing the importance of cognitive context in interpreting sensory phenotypes, and advocating for a shift beyond traditional sensory–cognitive dichotomies to better understand autism-related phenotypes.

### Learning alterations in autism and reliance on priors

Learning disability and intellectual disability are the second and third most common co-occurring neurodivergencies of autism, observed in 23.5% and 21.7% of autistic individuals, respectively (Khachadourian et al., 2023). Notably, even in the absence of these co-occurring conditions, autistic individuals exhibit differences in learning processes and altered neural mechanisms of learning (Mercado et al., 2020). In our study, we observed reduced performance in salience-based perceptual learning, shaped by a higher consistency bias in choice selection in *Fmr1*^-/y^ mice. These results align with findings in autistic individuals showing a stronger influence of previous choices on current perceptual decisions (Feigin et al., 2021). Moreover, decreased flexibility in perceptual decision-making is consistent with evidence of altered auditory and visual learning in autism (Harris et al., 2015; Alispahic et al., 2022), decreased performance under volatile conditions (Goris et al., 2021), as well as increased perseveration and diminished sensitivity to feedback (Crawley et al., 2020). Perceptual category learning was also found to be slower in autistic individuals, potentially due to the use of strategies that prioritize response guessing over rule application (Bott et al., 2006; Soulières et al., 2011). Impairments in visual, tactile, and audio-visual perceptual learning have also been shown in *Fmr1*^-/y^ mice (Arnett et al., 2014b; Goel et al., 2018; Mol et al., 2024).

Our findings extend this body of work by identifying a salience-dependent shift in reliance on priors in autism, with lower-salience stimuli more strongly evoking dependence on previous choices and impacting perceptual learning.

### Increased sensory discrimination in autism

Several theories of autism, including the *weak coherence* (Frith, 2003), *enhanced perceptual functioning* (Mottron et al., 2006), and *reduced generalization theories* (Plaisted, 2001), support the notion of a superior perception of low-level information in autism. Indeed, a body of work has confirmed superior pure tone discrimination (Bonnel et al., 2003; O’Riordan and Passetti, 2006) and discrimination learning of highly confusable patterns (Plaisted et al., 1998b). Tactile discrimination in autism has been studied using diverse experimental protocols that target different tactile features, which has often led to contradictory results due to methodological variability and participant heterogeneity (reviewed in (Zetler et al., 2019)). Similarly, work on roughness discrimination in rodent models (Orefice et al., 2016; Ahmadi et al., 2023) and human studies (O’Riordan and Passetti, 2006) has shown discrepancies.

To address translational challenges between animal models and clinical research, our approach employed vibrotactile stimuli closely aligned with those used in human studies. Consistent with findings in autistic individuals (He et al., 2021b), *Fmr1*^-/y^ mice showed intact amplitude discrimination for high-salience vibrotactile stimuli. Clinical studies have typically employed stimuli with more than 10-fold higher amplitudes compared to stimulus detection thresholds (He et al., 2021b; Asaridou et al., 2022). In contrast, our investigation focused on suprathreshold stimuli with much lower intensities (1.5-fold higher above threshold amplitude, 4-fold higher frequency). By employ such stimuli, our study uncovered enhanced discrimination of low-salience stimuli in *Fmr1*^-/y^ mice, an aspect that remains underexplored in autistic individuals. Increased “local” processing could account for enhanced discrimination performance (Frith, 2003). However, in our study, *Fmr1*^-/y^ mice exhibited enhanced discrimination only under specific stimulus salience conditions, arguing against a global increase in sensory processing precision. Future clinical studies assessing the discrimination of lower-level stimuli are necessary to explore whether enhanced low-salience discrimination also characterizes autistic individuals.

### Intact categorization but decreased facilitation of across-category discrimination in autism

Most perceptual discrimination is influenced by categorization (Goldstone, 1994; Beauny et al., 2020; Micher et al., 2024). As mentioned by Goldstone, “a clear distinction between sensory and cognitive processes is not tenable” (Goldstone, 1994). The reduced generalization hypothesis of autism postulates that, apart from superior performance on a difficult discrimination task, autistic individuals will have inferior performance in stimulus categorization (Plaisted, 2001). However, studies on rule-based and prototype categorization have yielded inconclusive results, reporting both intact (Klinger and Dawson, 2001; Molesworth et al., 2005), slower (Soulières et al., 2011), reduced (Froehlich et al., 2012; Gastgeb and Strauss, 2012) and enhanced performance (Bonnel et al., 2003) in categorization tasks in autistic individuals.

We assessed for the first time categorization and its impact on discrimination of vibrotactile stimuli in a mouse model of autism. Our results revealed intact tactile categorization but reduced facilitation of across-category discrimination in *Fmr1*^-/y^ mice. Although no clinical studies to date have examined this question in the tactile domain, our findings are consistent with previous research in the visual modality. These studies suggest that enhanced discrimination in autistic individuals may stem from reduced generalization (Plaisted et al., 1998b; Plaisted, 2001), and further indicate that discrimination processes may operate with increased independence from the influence of categorization (Soulières et al., 2007). These findings suggest a bidirectional interaction between sensory and cognitive alterations in autism, indicating that not only atypical sensory perception can impact cognition (Haigh, 2018), but also that cognitive differences can, in turn, shape sensory processing.

### Attentional alterations in autism

Although attention deficits are not considered a general characteristic of autism (Grubb et al., 2013), reduced attention in the presence of salient distractors (Venker et al., 2021) and weaker accuracy in executive attention (Ridderinkhof et al., 2018) have been reported in autistic individuals. Cognitive load is known to modulate attentional capacity (Lavie and Dalton, 2014; Murphy et al., 2016), and prior work has shown that high cognitive load alters selective attention and distractor filtering (Head and Helton, 2014; Lavie and Dalton, 2014; Brockhoff et al., 2022). However, less is known about attention to task-relevant stimuli in these conditions. Our results support the view that attention deficits emerge under conditions of high cognitive load in autism. Miss rates increased in our task when *Fmr^1^*^-/y^ mice were required to discriminate and categorize eight stimulus intensities within a single session for water reward, but not during training sessions with only two stimulus intensities.

Notably, cognitive load appears to disproportionately affect attention to low-salience stimuli in *Fmr1*^-/y^ mice, suggesting a disruption in late-stage attention processes, which take place after the stimulus is perceived and are involved in top-down goal-directed behavior (Calvillo and Jackson, 2014). In contrast, high-salience stimuli, which tend to capture attention automatically (Smallwood, 2013), may engage early attentional processes and thus less affected by cognitive load. This context-dependent attentional modulation aligns with previous findings in autistic individuals, showing stimulus-dependent auditory attention deficits (Čeponiene et al., 2003). These findings suggest that attentional alterations in autism may be conditionally expressed, particularly under tasks requiring increased cognitive resources.

Our results further dissociate attentional and tactile alterations in *Fmr1*^-/y^ mice, as low-salience stimuli were more accurately discriminated yet less attended to in *Fmr1*^-/y^ mice. These findings align with studies on linguistic processing in autistic children (Järvinen-Pasley et al., 2008; Ploog, 2010) and extend clinical research demonstrating that tactile sensitivity alterations in autism are not linked to attentional difficulties (He et al., 2021b).

### Trial history integration during perceptual decision-making in autism

Sensory perception is shaped not only by stimulus characteristics but also by trial history. Our results show that current stimulus intensity was the primary factor influencing the animal’s choice, together with a strong perseveration across genotypes, consistent with prior work on perceptual decision-making in mice (Aguillon-Rodriguez et al., 2021). This effect remained intact in *Fmr1*^-/y^ mice during tactile categorization. Interestingly, previous work in rats and humans has revealed that stimulus history strongly influences current perceptual choices under healthy conditions (Hachen et al., 2021). Our results confirmed these results in control mice and further revealed that this sensory history effect was absent in *Fmr1*^-/y^ mice. These findings suggest poor sensory history integration and support models of autism proposing faster world model updating, with new sensory information weighting higher than previous experience (Goris et al., 2022).

### Motivation and reward valuation during perceptual decision-making

Motivation and reward valuation are two crucial factors impacting performance during perceptual decision-making. In this study, we minimized confounds from fatigue, satiety, or disengagement by restricting analyses to sessions and epochs in which animals were actively performing the task. Under these conditions, *Fmr1*^-/y^ and WT mice showed indistinguishable task engagement, learning rates and learning trajectories, which would not be expected if motivation, reward value, or reinforcement efficacy were altered in our mouse model of autism.

During testing, only the sessions where the mice achieved high accuracy on well-learned training stimuli were included in the analysis. Several additional observations further argue against altered reward valuation as a primary driver of behavioral alterations in *Fmr1*^-/y^ mice. We observed no genotype differences in categorical response bias (criterion c), nor any effect of previous reward outcome on subsequent choices, indicating that reward history did not differentially influence decision-making in *Fmr1*^-/y^ mice. In contrast, we found reduced across-category discrimination facilitation and diminished influence of recent sensory history on choice, pointing to altered perceptual-decision processes rather than global motivational deficits. Moreover, *Fmr1*^-/y^ mice exhibited increased miss rates that emerged selectively under high cognitive load and reduced sensory salience conditions. The salience- and context-dependent nature of these effects, together with preserved engagement and reward sensitivity, supports the conclusion that the observed behavioral differences reflect specific disruptions in how sensory information is integrated and utilized during decision-making, rather than nonspecific changes in motivation, fatigue, or stereotyped responding.

### Conclusion and Future Perspectives

Numerous hypotheses have sought to provide a unified explanation for the diverse symptoms of autism. Our findings support the view that altered cognitive processes shape perception in autism, influencing how autistic individuals learn and engage with their environment. We demonstrate differences in choice bias during perceptual learning and increased tactile discrimination in trained animals. These changes are accompanied by reduced categorization influence, sensory history integration, and attention. Rather than reflecting a global sensory deficit or enhancement, our results point to context-dependent alterations in how sensory information is integrated, weighted, and used for decision-making.

Future work will be needed to identify the circuit mechanisms driving these effects, and to test whether they reflect intrinsic changes in top-down cognitive networks or altered feedback between cognitive and sensory circuits that reshape sensory representations. In addition, although the present study focused on amplitude discrimination within the flutter range (40 Hz) to align with translational tactile paradigms, extending this approach to a broader range of vibrotactile frequencies will be important to determine whether sensory–cognitive alterations in autism generalize across stimuli that engage different mechanoreceptor populations. Finally, while the present study focused on vibrotactile processing due to its strong translational relevance, previous studies in the visual modality have reported similar cognitive influences on perceptual decision-making, suggesting that these effects may generalize across sensory systems. Future studies directly comparing modalities within the same experimental framework will be important to confirm this. Elucidating how cognition modulates sensory perception may offer valuable insights for clinical practice, help reconcile conflicting findings in the field, and inform the development of more targeted, mechanism-based interventions.

## Materials and Methods

### Experimental design

To study tactile perception as well as attention and perceptual biases in autism, we developed a novel 2-Alternative Choice task for the categorization and discrimination of flutter-range vibrotactile stimuli. Throughout the text, we use terms that are preferred in the autistic community and are less stigmatizing (Bottema-Beutel et al., 2021).

### Ethical statement

All experimental procedures were performed in accordance with the EU directive 2010/63/EU and French law following procedures approved by the Bordeaux Ethics Committee and Ministry for Higher Education and Research. Mice were maintained in reversed light cycle under controlled conditions (temperature 22–24 °C, humidity 40–60%, 12 h/12 h light/dark cycle, light on at 21:00) in a conventional animal facility with *ad libitum* access to food and *ad libitum* access to water before the water restriction period. All experiments were performed during the dark cycle, under red light.

### Mice

Second-generation *Fmr1* knockout (*Fmr1*^−/y^) and wild-type littermate mice 5-16 weeks old were used in our study. Mice were maintained in a C57Bl/6 J background (Mientjes et al., 2006). Male wild-type and *Fmr1^-/y^* littermates were generated by crossing *Fmr1*^+/−^ females with *Fmr1*^+/y^ male mice from the same production, and the resulting progeny used for our experiments was either *Fmr1*^+/y^ (wild type) or *Fmr1^-/y^* (KO). Mice were maintained in collective cages following weaning (2-4 litter males per cage). Cages were balanced for genotype and supplemented with cotton nestlets and carton tubes.

The perceptual decision-making data were collected from 5 different cohorts of mice at different time-points during their active phase of the day. Mice of both genotypes were littermates and represented in each cohort. The number of mice is provided in the figure captions. The experimenter was blind to the animals’ genotypes throughout the experiment. The genotype of experimental animals was re-confirmed post hoc by tail-PCR.

### 2-Alternative Choice task Setup

The vibrotactile decision-making setup was positioned in an isolation cubicle to minimize interference during the experiment. Mice were placed in a body tube and were head-fixed with their forepaws resting on two steel bars (6 mm diameter, Thorlabs). The right bar was mounted to a Preloaded Piezo Actuator (P-841.6, Physik Instrumente) equipped with a strain gauge feedback sensor and controlled (E-501, Physik Instrumente) in a closed loop, as described before (Prsa et al., 2019; Semelidou et al., 2024). A 12.7 mm stainless steel post (ThorLabs) was mounted on the actuator vertically and a 0.6 mm stainless steel rod (ThorLabs) was clamped horizontally onto this post. The horizontal rod served as the contact bar on which the animal rested its right forepaw. Water reward was delivered through either of the two metal feeding needles (20G, 1,9mm tip, Agntho’s AB), placed left and right of the mouse’s mouth, each connected to a lickport interface with a solenoid valve (Sanworks) equipped with a capacitive sensor (https://github.com/poulet-lab/Bpod_CapacitivePortInterface). The perceptual decision-making setup was controlled by Bpod (Sanworks) through scripts in Python (PyBpod, https://pybpod.readthedocs.io/en/latest/). The lickport interface (Sanworks) was equipped with a capacitive sensor (https://github.com/poulet-lab/Bpod_CapacitivePortInterface).

### Habituation to head-fixation and water restriction

Mice (P40-P50) were handled using carton tubes and the cupping technique until they were comfortable in the experimenter’s hands, attested by eating while handled. Mice were gradually habituated to the experimental setup and head fixation for 5 days. The third day of habituation, a water-restriction protocol was implemented, where mice had access to liquid water in the setup and to a solid water supplement (Hydrogel, BioServices) in their home cage. The water supplement was divided into small, individual portions, and each mouse received its allotment after the daily training/testing session. Daily body weight measurements were used to monitor hydration and ensure that all animals maintained stable body weight. If necessary, supplemental water was adjusted to maintain animals within the approved weight range. In total, the animals received 1.5-2 ml of water per day, which corresponds to 50-65% of their ad libitum consumption, while ensuring that they did not lose more than 10% of their weight. Each mouse received 6-8 g of Hydrogel (*ad libitum*) during the weekend. This water restriction protocol was maintained throughout behavioral training and until the end of behavioral testing.

### 2-Alternative Choice task training and testing Vibrotactile Stimuli

Stimuli were sinusoidal vibrations at 40 Hz with peak-to-peak displacements of 12 μm amplitude (low salience) or 26 μm amplitude (high salience), well above the detection threshold of both groups (10 Hz, 4.46 µm for WT; 10Hz, 7.29 µm for *Fmr1*^-/y^ mice) (Semelidou et al., 2024).

Animals were positioned in the setup to ensure stable and consistent forepaw contact with the rod delivering the vibration. Pilot experiments with a sensor to monitor forepaw placement confirmed that the mice did not remove their forepaws from the bar before stimulus delivery.

Analysis of trials were no response followed stimulus delivery (Miss trials) showed that the 12 μm stimulus consistently elicited a higher proportion of missed responses compared to the 26 μm stimulus across animals, indicating lower behavioral performance for the lower-amplitude stimulus. We therefore referred to the 12 μm stimulus as “low salience” and the 26 μm stimulus as “high salience” to denote relative differences in perceptual strength and attentional engagement within the suprathreshold range.

Habituated mice (8 weeks old) were trained to associate high- (26 μm) and low-salience (12 μm) vibrotactile stimuli (pure sinusoid, 500 ms duration, 40 Hz frequency) with a water reward (8 µl) at the right- or left-placed lickport of the setup, respectively. All trials consisted of stimulus delivery followed by a 2 s response window during which the mice could lick to receive the reward. Inter-trial intervals were variable (5-10 s). Training was subdivided in 4 phases: (a) automatic water delivery at the beginning of the response window at the corresponding lickport (left for 12 μm stimuli, right for 26 μm stimuli). (b) Training in blocks: lick-triggered water delivery during blocks of 20 trials with the same vibrotactile stimulus, of either 12 μm or 26 μm amplitude. Licking at the wrong port resulted in 5 s timeout. (c) Training with pseudorandomly delivered high- and low-salience trials: lick-triggered water delivery following pseudorandom delivery of 12 μm or 26 μm amplitude stimuli. Licking at the wrong port resulted in 5 s timeout.

All sessions consisted of 200-300 trials with a 1:1 ratio of high and low-salience stimuli. During training, this ratio was modified when an animal showed consistent bias for one of the two lickports. Pilot experiments with an extra sensor to monitor forepaw placement confirmed that the mice did not remove their forepaws from the bar before stimulus delivery. To complete training in blocks, mice needed to perform with 70% correct choices and 30% incorrect choices for both stimulus amplitudes. To complete training with pseudorandomly delivered high- and low-salience trials, mice needed to reach the criterion of more than 70% correct choices and less than 30% incorrect choices as an average for 3 consecutive days. All mice that fulfilled this criterion were tested for the categorization/discrimination of vibrotactile stimuli. During testing, stimuli (pure sinusoid, 500 ms duration, 40 Hz frequency) were delivered in a pseudorandom manner with a 50% High-salience: 50% Low-salience ratio. Amplitudes varied on a range between 12 μm and 26 μm. Stimuli of 20, 22, 24 and 26 μm were considered high-salience stimuli and rewarded at the right lickport while stimuli of 12, 14, 16 and 18 μm were considered as low-salience stimuli and rewarded at the left lickport.

### 2-Alternative choice vibrotactile task analysis

All analysis was performed with custom-made Python scripts that can be available upon request. Behavior was quantified based on the lick events and three main outcomes were measured for each stimulus salience: Correct choice rate (number of correct licks divided by the total number of high- or low-salience trials), Incorrect choice rate (number of wrong licks divided by the total number of high- or low-salience

trials), and Missed stimuli rate (number of trials in which the animal did not lick, divided by the total number of high- or low-salience trials). Sessions in which animals disengaged were analyzed only during epochs in which the animal was actively performing the task. Training duration was calculated based on the total number of days each animal passed in training in blocks and with pseudorandom stimulus delivery.

For testing, only sessions with more than 70% correct choices for the training stimuli (12 μm and 26 μm) were analyzed. Psychometric curves were fitted on the rightwards lick rate for each stimulus amplitude using a general linear model. An average of 84 repetitions for each amplitude was used to calculate rightward lick rates.

Based on the signal detection theory (Detection Theory,(Hautus, M.J., Macmillan, N.A., & Creelman, 2021)), sensitivity d’ was calculated as:

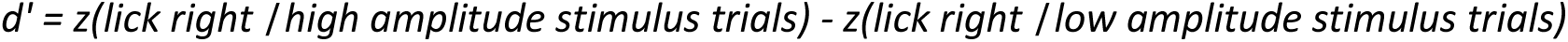

Sensitivity d’ for high- and low-salience stimuli was calculated based on the Correct and Incorrect choice rate for high- and low-salience stimuli, respectively.

The strategy of the animals was assessed through their criterion, which was calculated as:

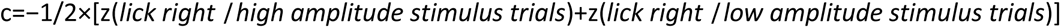

Categorization bias was calculated as the inflection point (i.e., the midpoint parameter μ) of the fitted logistic psychometric function and categorization precision as the slope of the curve.

Prior choice impact was calculated separately for each type of the current trial (high- or low-salience). For low-salience trials, the prior choice impact was calculated as the rate of correct low and incorrect-low trials, divided by the rate of all correct and all incorrect trials. Similarly, for high-salience trials, the prior choice impact was calculated as the rate of correct high and incorrect-high trials, divided by the rate of all correct and all incorrect trials.

Learning trajectory analysis

Learning curves were quantified from the training phase with mixed high- and low-salience trials by calculating, for each mouse, the proportion of correct choices on each training day. For group-level comparisons, performance was plotted across days for WT and *Fmr1*^-/y^ mice, and mean learning trajectories with 95% confidence intervals were computed for each genotype.

To statistically compare learning trajectories across genotypes, we used a linear mixed-effects model with fixed effects for Genotype, Day, and their interaction, and a random intercept and random slope for each animal. This model tested whether performance differed between genotypes across days and whether the rate of learning (performance change per day) differed between WT and *Fmr1*^-/y^ mice.

To quantify individual learning rates, we computed the slope of performance as a function of training day for each animal. For each mouse, performance values (proportion of correct choices) were regressed against each Day using a simple linear regression model:

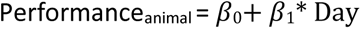

The slope (β₁) provides an estimate of the rate of improvement per training day.

Intermediate-stage slopes (middle 3 days of each mouse’s training) were calculated in the same manner. Categorization bias was calculated based on the psychometric curves, as the stimulus amplitude at the inflection point of the sigmoid fitting curve. Categorization precision was calculated based on the slope of the psychometric curve.

Delta discrimination accuracy was computed as the difference in the rate of high-salience (rightward) licks between pairs of stimuli that differed by 2 µm. For analyses of low-salience and high-salience discrimination, delta discrimination accuracy values were calculated for each within-category stimulus pair and then averaged within the low-salience or high-salience category, respectively.

Statistics

All values are presented as mean ± s.e.m. Box plots show the median, interquartile, range, mean and individual values. The numbers of animals used are indicated in the figure legends. Sample size was determined based on previous studies using similar behavioral paradigms. For experiments assessing tactile responses of trained animals, only animals that reached the training criterion were tested. For the discrimination & categorization task, all the testing sessions where the animals showed less than 70% correct responses to the training stimuli (12µm and 26µm) were excluded from tactile discrimination/categorization analysis. No samples or data points were excluded from the analysis.

All statistical analysis was done using Python (SciPy, Pingouin). All datasets were tested for normality using Shapiro–Wilk tests, and potential outliers were identified using a median absolute deviation–based modified Z-score (|Z| > 3.5). When an outlier was detected, statistical analyses were performed both including and excluding the outlier. In such cases, results from both analyses are reported when the inclusion or exclusion of the outlier altered the statistical outcome, to ensure transparency and robustness of the findings. For normally distributed data, two-tailed paired t-tests were used for within-subject comparisons, and Welch’s independent-samples t-tests were used for between-group comparisons; otherwise, the Mann–Whitney U test (between groups) or the Wilcoxon signed-rank test (within subjects) was applied. Bonferroni correction was applied where multiple comparisons were performed. Effect sizes were calculated using Hedge’s g for between-genotype comparisons and paired Hedge’s g for within-genotype comparisons, particularly when statistical analyses showed trend-level differences (p ≈ 0.05–0.1), to provide a measure of the magnitude of the effect independent of sample size. Linear mixed-effects models were used to assess statistical differences in learning trajectories, psychometric curves, and miss rates.

Statistical modeling during categorization taking into account the trial history. Generalized linear models (GLMs) with a binomial link function were used to quantify the effects of current stimulus, previous trial history, genotype, and their interactions on the probability of choosing the right port. Trials without a defined history and Miss trials were excluded. Stimulus amplitudes were z-scored per mouse.

Two models were fit:

1. Main effects GLM – included current stimulus, previous stimulus, previous outcome, previous choice, and genotype.
2. Interaction GLM – extended the main effects model by including genotype × previous outcome, genotype × previous choice, and genotype × previous stimulus interactions.

Cluster-robust standard errors were calculated to account for repeated measurements within each mouse.

Marginal effects and psychometric curves

Predicted probabilities were computed from the interaction GLM to visualize the effect of stimulus amplitude, previous choice, and previous outcome on behavior, separately for WT and KO mice.

## Data availability

The raw data of behavioral experiments generated in this study have been deposited on the figshare database: 10.6084/m9.figshare.29459771. Source data for each figure and the Python code used for all analyses are provided with the paper.

## Figures and Figure Legends

**Figure S1.**
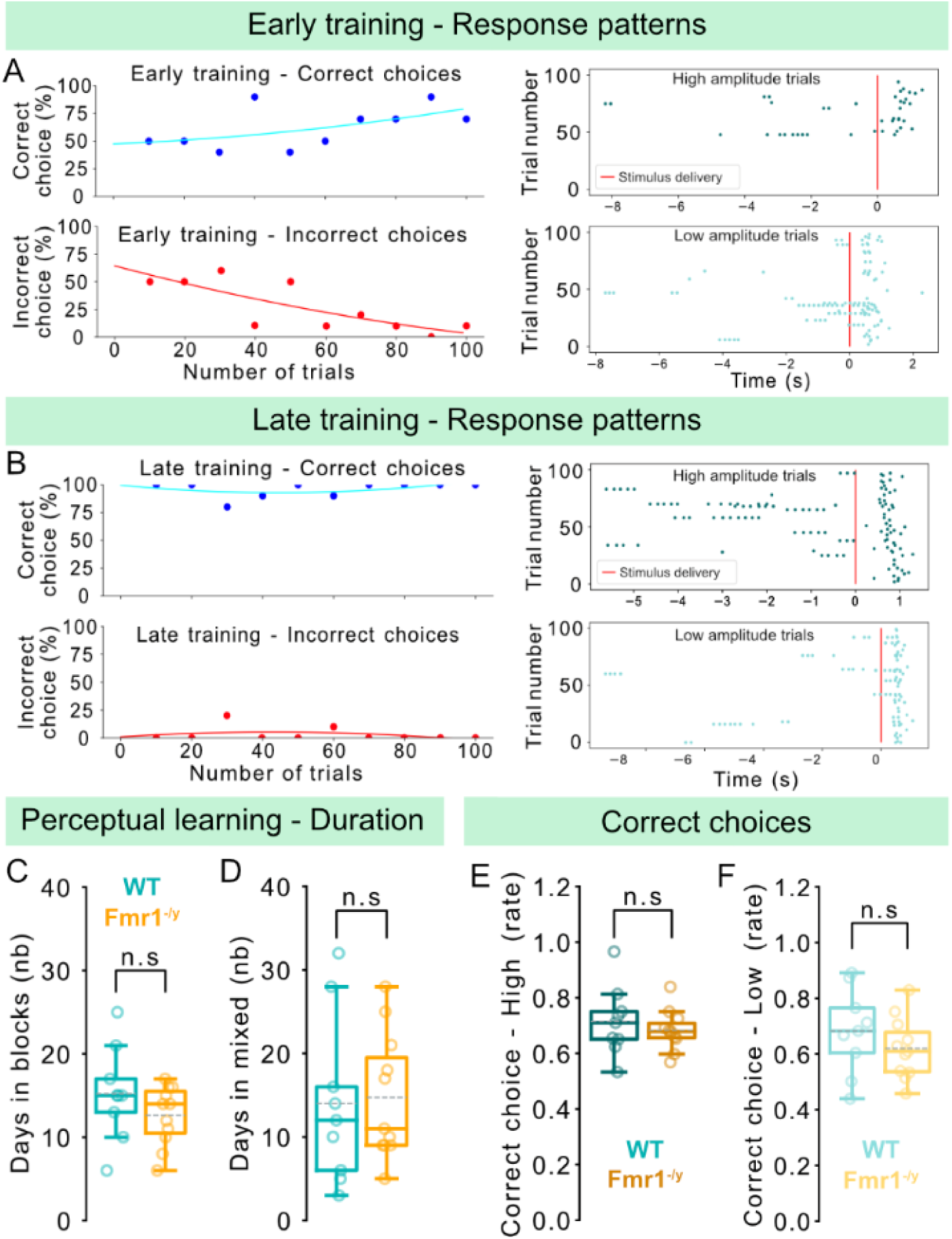
Perceptual learning duration for the different training phases and correct choice rates. n=9 WT, 11 *Fmr1*^-/y^ male mice for panels **C-F**. **A,** (Left) Example trace of 100 consecutive trials from an early (mixed) training session, showing the rates of correct (top) and incorrect (bottom) choices. (Right) Licking patterns from the same session for high-salience (top) and low-salience (bottom) trials. **B,** Same as in (A), but from a late training session of the same animal. **C,** Total number of days spent in the training phase where stimulus delivery is done in blocks of high- or low-salience stimuli. **D,** Total number of days spent in training with high-and low-salience stimuli are delivered in a pseudorandom manner. **E,** Correct choice rate for high-salience trails throughout the training period for WT and *Fmr1*^-/y^ mice. **F,** Correct choice rate for low-salience trails throughout the training period for WT and *Fmr1*^-/y^ mice. P values were computed using two-sided t-test for panels **C, D, E, F**; n.s, not significant.

**Figure S2.**
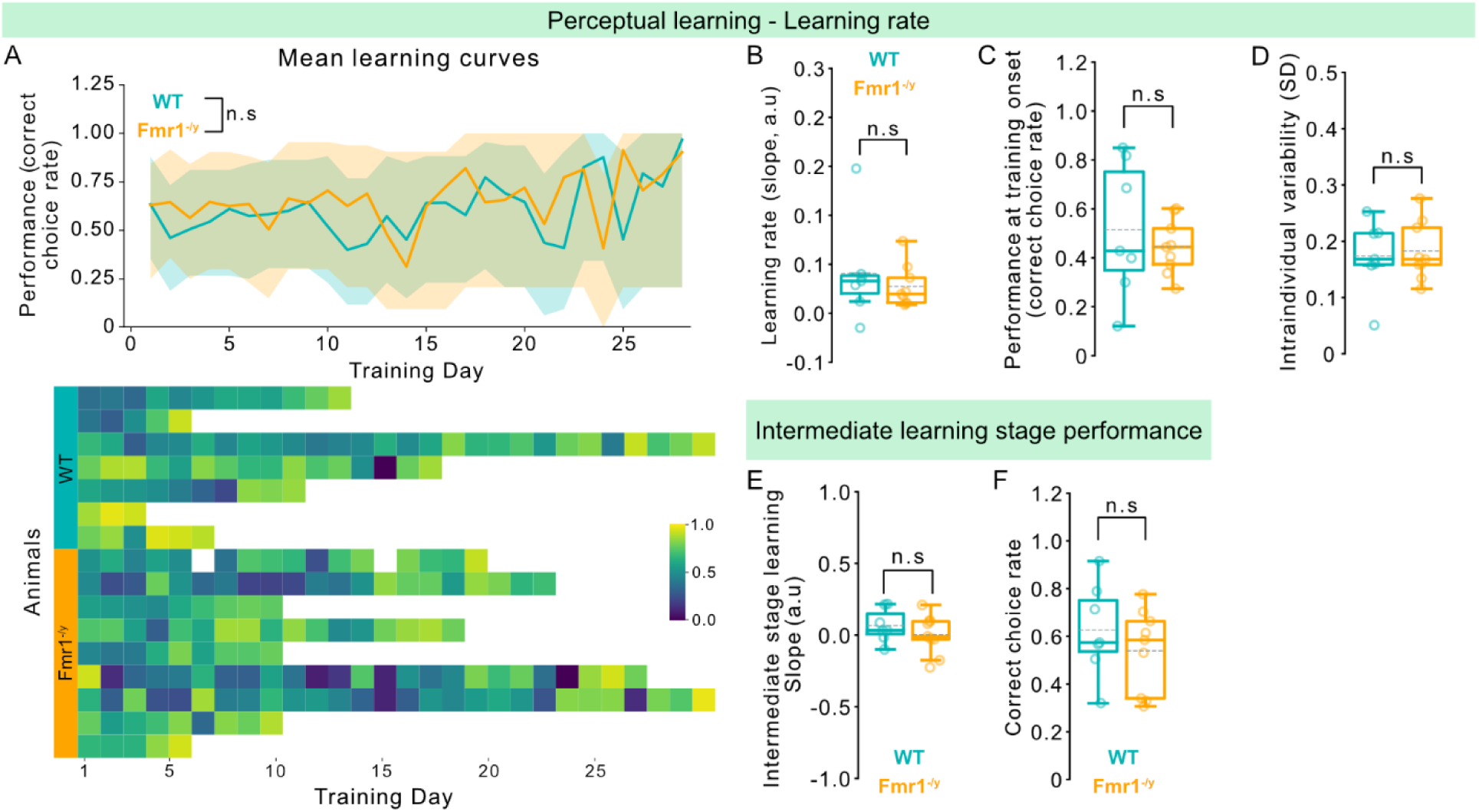
Perceptual learning trajectories during training with mixed low- and high-salience trials. n=7 WT, 9 *Fmr1*^-/y^ male mice for panels. **A,** (Top) Average learning trajectories for WT and *Fmr1*^-/y^ mice that completed the training phase, during mixed-trial training, calculated from the correct-choice rate for each training session (day) until each mouse reached the learning criterion. Shaded areas indicate 95% confidence intervals. (Bottom) Heat map of individual performance across mixed high- and low-salience sessions, showing the correct-choice rate for each session up to each mouse’s learning criterion. **B,** Slope of the learning trajectory for each mouse, calculated using a linear regression model. **C,** Performance (correct choice rate) on the first day of the mixed-trial training phase. **D,** Intra-individual variability in performance across training days, assessed as the standard deviation of the correct choice rate. **E,** Slope of the learning trajectory during the intermediate learning stage, defined as the middle 3 days of training for each animal. **F,** Performance (correct choice rate) during the intermediate learning stage. P values were computed using linear mixed-effects model for panel **A**, Mann-Whitney test for panel **B**, and two-sided t-test for panels **C, D, E, F**; n.s, not significant.

**Figure S3.**
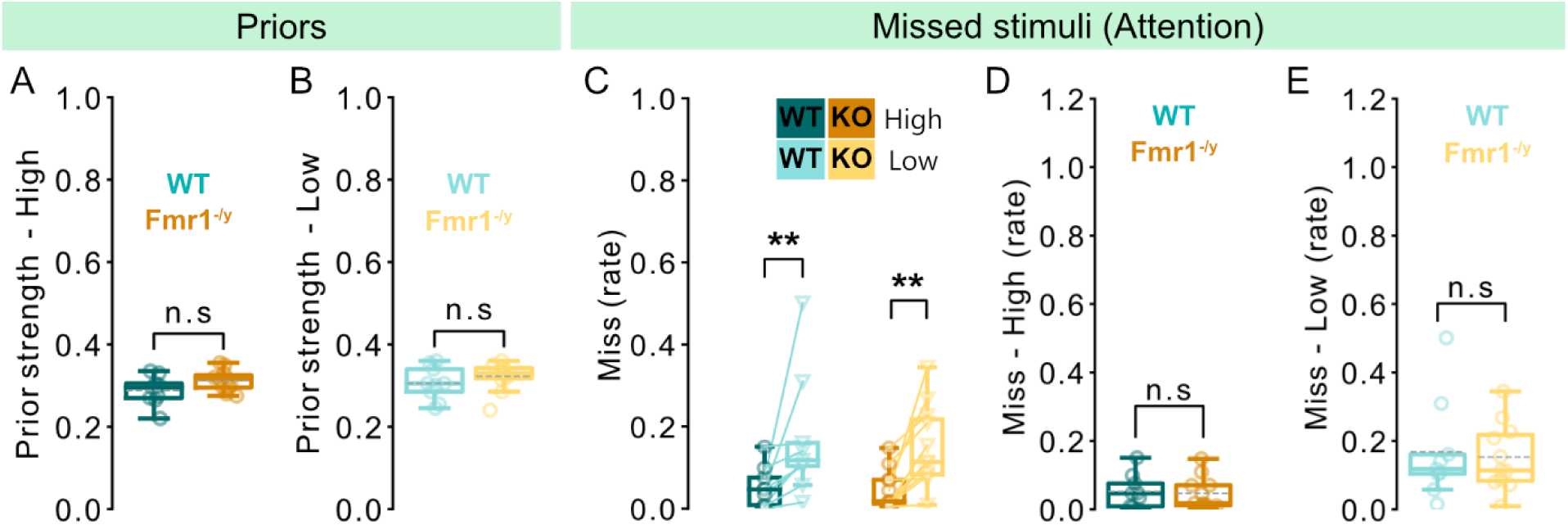
Prior strength and attention during perceptual learning. n=9 WT, 11 *Fmr1*^-/y^ male mice. **A,** Strength of the prior build for high-salience trials, calculated as the proportion of correct high-salience and incorrect low-salience responses following a correct high-salience response. Rates are corrected over the rate of overall correct high-salience and incorrect low-salience responses. **B,** Strength of the prior build for low-salience trials, calculated as the proportion of correct low-salience and incorrect high-salience responses following a correct low-salience response. Rates are corrected over the rate of overall correct low-salience and incorrect high-salience responses. **C,** Within-genotype comparisons of the proportion of missed high- and low-salience trials. **D,** Proportion of missed high-salience trials. **E,** Proportion of missed low-salience trials. P values were computed using two-sided t-test for panel **A,**; Mann-Whitney test for panels **B, D, E**; Wilcoxon signed-rank test for panel **C-left;** Paired t-test for panel **C-right**,; **P < 0.01, or n.s, not significant.

**Figure S4.**
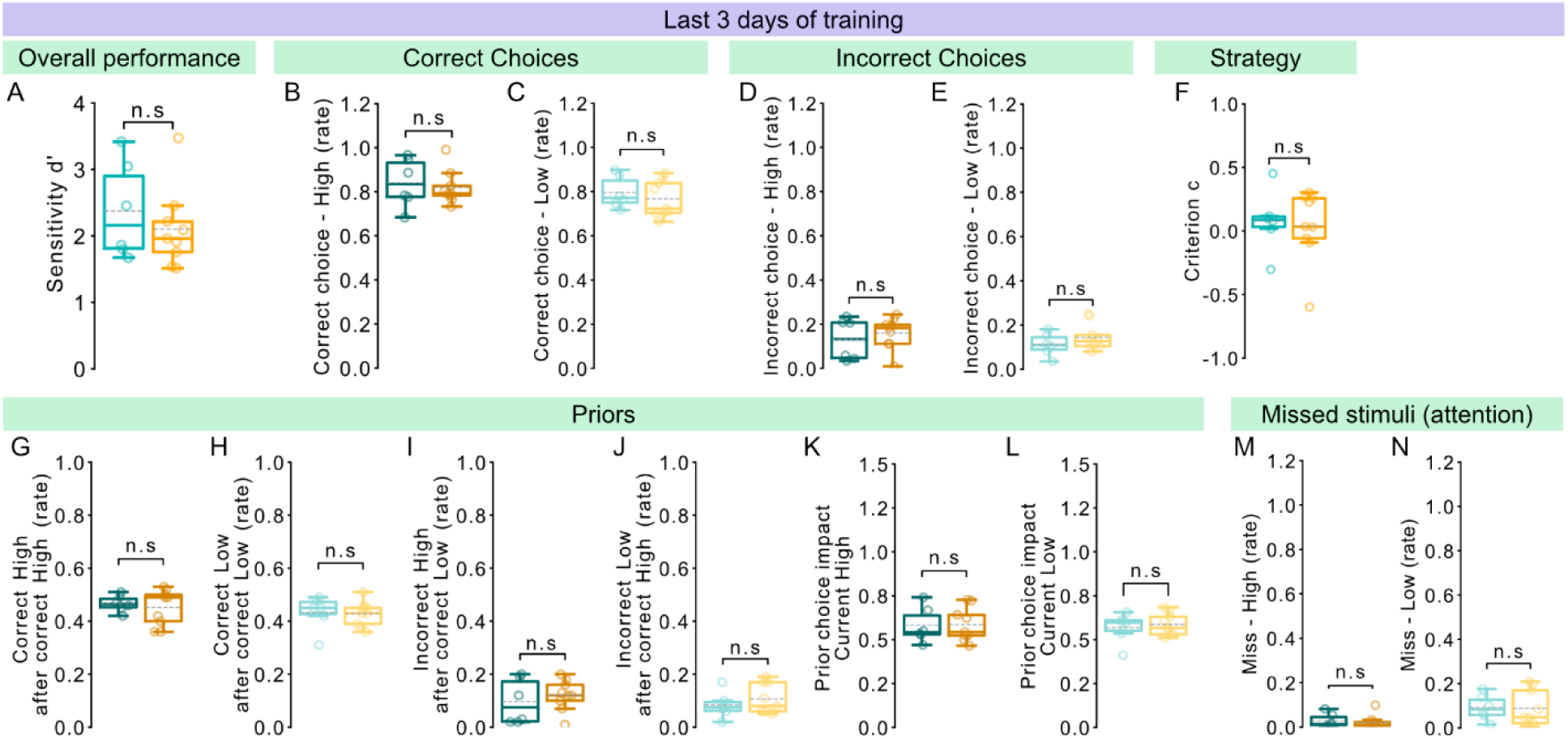
Perceptual learning performance during the last three days of training, for mice that were tested in tactile discrimination. n=6 WT, 9 *Fmr1*^-/y^ male mice. **A,** Sensitivity d’ throughout the last three days of the training period for WT and *Fmr1*^-/y^ mice. **B,** Proportion of correct choices for high-salience trails throughout the last three days of the training period. **C,** Proportion of correct choices for low-salience trails throughout the last three days of the training period. **D,** Proportion of incorrect choices for high-salience trails throughout the last three days of the training period. **E,** Proportion of incorrect choices for low-salience trails throughout the last three days of the training period. **F,** Criterion depicting the licking strategy of the animals. **G,** Proportion of correct responses in high-salience trials immediately following a correctly rewarded high-salience trial. **H,** Proportion of correct responses in low-salience trials immediately following a correctly rewarded low-salience trial. **I,** Proportion of incorrect responses in high-salience trials immediately following a correctly rewarded low-salience trial. **J,** Proportion of incorrect responses in low-salience trials immediately following a correctly rewarded high-salience trial. **K,** Proportion of correct responses in high-salience trials immediately following a correctly rewarded high-salience trial and incorrect responses in high-salience trials immediately following a correctly rewarded low-salience trial. Rates are corrected over the total number of correct and incorrect choices in high-salience trials. **L,** Proportion of correct responses in low-salience trials immediately following a correctly rewarded low-salience trial and incorrect responses in low-salience trials immediately following a correctly rewarded high-salience trial. Rates are corrected over the total number of correct and incorrect choices in low-salience trials. **M,** Proportion of missed high-salience trials. **N,** Proportion of missed low-salience trials. P values were computed using two-sided t-test for panels **A, B, C, D, E, G, H, I, J, K, L, N**; Mann-Whitney test for panel **F**, **M,**; n.s, not significant.

**Figure S5.**
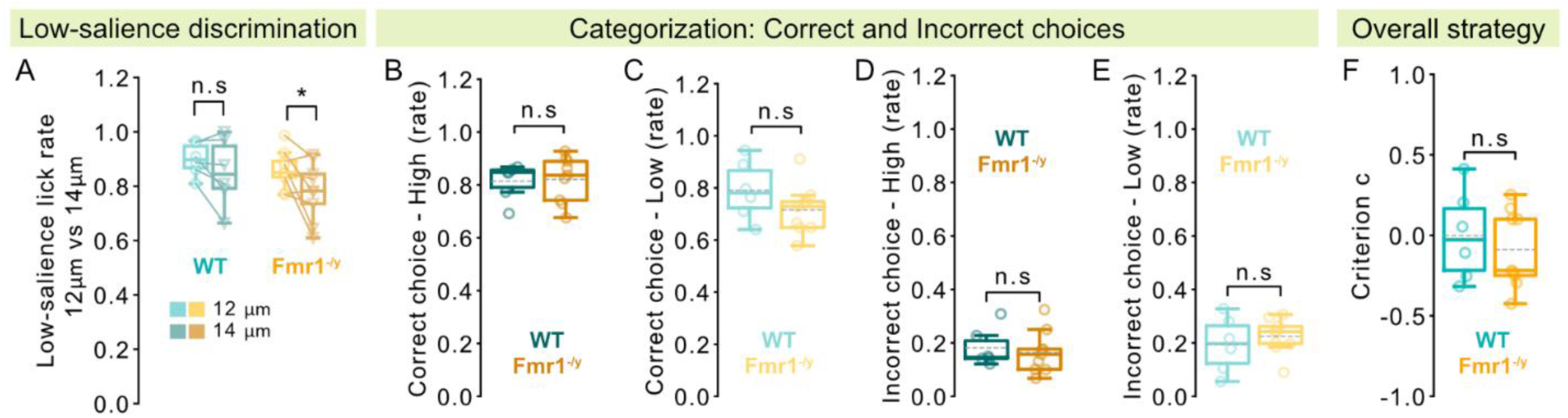
Sensitivity, correct, and incorrect choices during categorization of high- and low-salience stimuli. n=6 WT, 9 *Fmr1*^-/y^ male mice. **A,** Sensitivity d’ for high-salience stimuli during categorization. **B,** Sensitivity d’ for low-salience stimuli during categorization. **C,** Correct responses for high-salience stimuli during categorization. **D,** Correct responses for low-salience stimuli during categorization. **E,** Incorrect responses for high-salience stimuli during categorization. **F,** Incorrect responses for low-salience stimuli during categorization. P values were computed using two-sided paired t-test for panel **A,**; two-sided t-test for panels **B, C, D, E, F,;** *P < 0.05, or n.s, not significant.

## Acknowledgments

We would like to thank the animal housing and genotyping facilities of INSERM U1215 Neurocenter Magendie. We thank Dr. James Poulet (Max-Delbrück-Centrum für Molekulare Medizin, Berlin) for providing the capacitive sensors for the behavioral task. We would like to thank Dr. Melanie Ginger for providing feedback on the manuscript. Mouse schemas were made using Biorender. This project was funded by the Foundation for Medical Research Postdoc Fellowship (O.S.), INSERM (A.F.), Marcel Dassault-Fondation FondaMental Award 2019 (A.F.), Simons Foundation Autism Research Initiative (A.F.), GIS Autisme & TND (O.S.), 2024 NARSAD Young Investigator Grant (O.S.), Fondation FondaMental (A.F.).

## Author Information Authors and Affiliations

*University of Bordeaux, Neurocentre Magendie, INSERM U1215, 33000, Bordeaux, France* Ourania Semelidou, Mathilde Tortochot-Megne Fotso, Adinda Winderickx, Andreas Frick

## Contributions

A.F. and O.S. conceived the project. O.S. designed the experiments. O.S., M.T-M.F. and A.W. performed the experiments. O.S. and A.W. wrote the Python code for the behavioral experiments. O.S. analyzed the data and wrote the necessary Python scripts. O.S. interpreted the data. O.S. prepared the figures. O.S. wrote the manuscript, and A.F. and M.T-M.F. provided feedback.

## Corresponding author

Correspondence to Andreas Frick and Ourania Semelidou.

## Ethics declarations

Competing interests

The authors declare no competing interests.

## Inclusion and diversity statement

We support inclusive, diverse, and equitable conduct of research. We tried to use inclusive language as much as possible.

## References

Abu-Dahab SMN, Skidmore ER, Holm MB, Rogers JC, Minshew NJ (2013) Motor and Tactile-Perceptual Skill Differences Between Individuals with High-Functioning Autism and Typically Developing Individuals Ages 5 – 21. J Autism Dev Disord 43:2241 Available at: https://pmc.ncbi.nlm.nih.gov/articles/PMC3408783/ [Accessed April 16, 2025].

Aguillon-Rodriguez V et al. (2021) Standardized and reproducible measurement of decision-making in mice. Elife 10 Available at: /pmc/articles/PMC8137147/ [Accessed November 23, 2022].

Ahmadi BBM, Afarinesh MR, Jafaripour L, Sheibani V (2023) Alteration in social interaction and tactile discrimination of juvenile autistic-like rats following tactile stimulation and whisker deprivation. Brain Behav 13.

Alispahic S, Pellicano E, Cutler A, Antoniou M (2022) Auditory perceptual learning in autistic adults. Allen G, Courchesne E (2001) Attention function and dysfunction in autism. Front Biosci 6 Available at: https://pubmed.ncbi.nlm.nih.gov/11171544/ [Accessed June 23, 2025].

American Psychiatric Association (2022) Diagnostic and statistical manual of mental disorders (5th ed., text rev.). American Psychiatric Association Publishing. Available at: https://psychiatryonline.org/doi/book/10.1176/appi.books.9780890425787.

Amoruso L, Narzisi A, Pinzino M, Finisguerra A, Billeci L, Calderoni S, Fabbro F, Muratori F, Volzone A, Urgesi C (2019) Contextual priors do not modulate action prediction in children with autism. Proc R Soc B 286 Available at: https://royalsocietypublishing.org/doi/10.1098/rspb.2019.1319 [Accessed April 23, 2023].

Arnett MT, Herman DH, McGee AW (2014a) Deficits in Tactile Learning in a Mouse Model of Fragile X Syndrome. PLoS One 9 Available at: /pmc/articles/PMC4189789/ [Accessed May 9, 2024].

Arnett MT, Herman DH, McGee AW (2014b) Deficits in tactile learning in a mouse model of fragile X syndrome. PLoS One 9 Available at: https://pubmed.ncbi.nlm.nih.gov/25296296/ [Accessed May 28, 2024].

Asaridou M, Wodka EL, Edden RAE, Mostofsky SH, Puts NAJ, He JL (2022) Could Sensory Differences Be a Sex-Indifferent Biomarker of Autism? Early Investigation Comparing Tactile Sensitivity Between Autistic Males and Females. J Autism Dev Disord:1–17 Available at: https://link.springer.com/article/10.1007/s10803-022-05787-6 [Accessed December 11, 2022].

Balasco L, Pagani M, Pangrazzi L, Chelini G, Ciancone Chama AG, Shlosman E, Mattioni L, Galbusera A, Iurilli G, Provenzano G, Gozzi A, Bozzi Y (2021) Abnormal Whisker-Dependent Behaviors and Altered Cortico-Hippocampal Connectivity in Shank3b −/− Mice. Cereb Cortex:1–14.

Balasco L, Pagani M, Pangrazzi L, Chelini G, Viscido F, Chama AGC, Galbusera A, Provenzano G, Gozzi A, Bozzi Y (2022) Somatosensory cortex hyperconnectivity and impaired whisker-dependent responses in Cntnap2 -/- mice. Neurobiol Dis 169 Available at: https://pubmed.ncbi.nlm.nih.gov/35483565/ [Accessed July 19, 2022].

Barbot A, Liu S, Kimchi R, Carrasco M (2018) Attention enhances apparent perceptual organization. Psychon Bull Rev 25:1824 Available at: https://pmc.ncbi.nlm.nih.gov/articles/PMC5831484/ [Accessed May 20, 2025].

Beauny A, de Heering A, Muñoz Moldes S, Martin JR, de Beir A, Cleeremans A (2020) Unconscious categorization of submillisecond complex images. PLoS One 15 Available at: https://pubmed.ncbi.nlm.nih.gov/32785238/ [Accessed May 23, 2025].

Bonnel A, Mottron L, Peretz I, Trudel M, Gallun E, Bonnel AM (2003) Enhanced pitch sensitivity in individuals with autism: A signal detection analysis. J Cogn Neurosci 15:226–235 Available at: https://pubmed.ncbi.nlm.nih.gov/12676060/ [Accessed May 22, 2025].

Bott L, Brock J, Brockdorff N, Boucher J, Lamberts K (2006) Perceptual similarity in autism. Q J Exp Psychol 59:1237–1254 Available at: https://pubmed.ncbi.nlm.nih.gov/16769623/ [Accessed April 26, 2025].

Bottema-Beutel K, Kapp SK, Lester JN, Sasson NJ, Hand BN (2021) Avoiding Ableist Language: Suggestions for Autism Researchers. Autism adulthood challenges Manag 3:18–29 Available at: https://pubmed.ncbi.nlm.nih.gov/36601265/ [Accessed July 17, 2024].

Bouvier B, Susini P, Marquis-Favre C, Misdariis N (2023) Revealing the stimulus-driven component of attention through modulations of auditory salience by timbre attributes. Sci Rep 13:1–11 Available at: https://www.nature.com/articles/s41598-023-33496-2 [Accessed April 30, 2025].

Bremner AJ, Spence C (2017) The Development of Tactile Perception. Adv Child Dev Behav 52:227–268.

Brockhoff L, Schindler S, Bruchmann M, Straube T (2022) Effects of perceptual and working memory load on brain responses to task-irrelevant stimuli: Review and implications for future research. Neurosci Biobehav Rev 135:104580 Available at: https://www.sciencedirect.com/science/article/abs/pii/S0149763422000690 [Accessed May 23, 2025].

Calvillo DP, Jackson RE (2014) Animacy, perceptual load, and inattentional blindness. Psychon Bull Rev 21:670–675 Available at: https://link.springer.com/article/10.3758/s13423-013-0543-8 [Accessed May 7, 2025].

Carati E, Angotti M, Pignataro V, Grossi E, Parmeggiani A (2024) Exploring sensory alterations and repetitive behaviors in children with autism spectrum disorder from the perspective of artificial neural networks. Res Dev Disabil 155 Available at: https://pubmed.ncbi.nlm.nih.gov/39577022/ [Accessed November 26, 2024].

Cascio CJ, Gu C, Schauder KB, Key AP, Yoder P (2015) Somatosensory event-related potentials and association with tactile behavioral responsiveness patterns in children with ASD. Brain Topogr 28:895 Available at: /pmc/articles/PMC4601930/ [Accessed December 3, 2022].

Čeponiene R, Lepistö T, Shestakova A, Vanhala R, Alku P, Näätänen R, Yaguchi K (2003) Speech-sound-selective auditory impairment in children with autism: They can perceive but do not attend. Proc Natl Acad Sci U S A 100:5567–5572 Available at: /doi/pdf/10.1073/pnas.0835631100?download=true [Accessed May 6, 2025].

Church B, Krauss MS, Lopata C, Toomey JA, Thomeer ML, Coutinho M V., Volker MA, Mercado E (2010) Atypical categorization in children with high-functioning autism spectrum disorder. Psychon Bull Rev 17:862–868.

Crawley D, Zhang L, Jones EJH, Ahmad J, Oakley B, Cáceres ASJ, Charman T, Buitelaar JK, Murphy DGM, Chatham C, den Ouden H, Loth E (2020) Modeling flexible behavior in childhood to adulthood shows age-dependent learning mechanisms and less optimal learning in autism in each age group. PLoS Biol 18:e3000908 Available at: https://pmc.ncbi.nlm.nih.gov/articles/PMC7591042/ [Accessed April 26, 2025].

Demopoulos C, Brandes-Aitken AN, Desai SS, Hill SS, Antovich AD, Harris J, Marco EJ (2015) Shared and Divergent Auditory and Tactile Processing in Children with Autism and Children with Sensory Processing Dysfunction Relative to Typically Developing Peers. J Int Neuropsychol Soc 21:444–454 Available at: https://www.cambridge.org/core/journals/journal-of-the-international-neuropsychological-society/article/abs/shared-and-divergent-auditory-and-tactile-processing-in-children-with-autism-and-children-with-sensory-processing-dysfunction-relative-to-typically-developing-peers/0F61C695B7ED89D61F5150C0978F209B [Accessed October 12, 2023].

Espenhahn S, Godfrey KJ, Kaur S, McMorris C, Murias K, Tommerdahl M, Bray S, Harris AD (2023) Atypical Tactile Perception in Early Childhood Autism. J Autism Dev Disord 53:2891–2904 Available at: https://pubmed.ncbi.nlm.nih.gov/35482274/ [Accessed October 12, 2023].

Failla MD, Peters BR, Karbasforoushan H, Foss-Feig JH, Schauder KB, Heflin BH, Cascio CJ (2017) Intrainsular connectivity and somatosensory responsiveness in young children with ASD. Mol Autism 8:1–11 Available at: https://molecularautism.biomedcentral.com/articles/10.1186/s13229-017-0143-y [Accessed April 16, 2025].

Feigin H, Shalom-Sperber S, Zachor DA, Zaidel A (2021) Increased influence of prior choices on perceptual decisions in autism. Elife 10.

Forschack N, Gundlach C, Hillyard S, Müller MM (2022) Attentional capture is modulated by stimulus saliency in visual search as evidenced by event-related potentials and alpha oscillations. Attention, Perception, Psychophys 2022 853 85:685–704 Available at: https://link.springer.com/article/10.3758/s13414-022-02629-6 [Accessed April 30, 2025].

Foss-Feig JH, Heacock JL, Cascio CJ (2012) Tactile responsiveness patterns and their association with core features in autism spectrum disorders. Res Autism Spectr Disord 6:337–344.

Frith U (2003) Autism: Explaining the enigma, 2nd ed. Malden: Blackwell Publishing.

Froehlich AL, Anderson JS, Bigler ED, Miller JS, Lange NT, Dubray MB, Cooperrider JR, Cariello A, Nielsen JA, Lainhart JE (2012) Intact Prototype Formation but Impaired Generalization in Autism. Res Autism Spectr Disord 6:921 Available at: https://pmc.ncbi.nlm.nih.gov/articles/PMC3267426/ [Accessed May 22, 2025].

Gastgeb HZ, Strauss MS (2012) Categorization in ASD: The Role of Typicality and Development. Perspect Lang Learn Educ 19:66 Available at: https://pmc.ncbi.nlm.nih.gov/articles/PMC3372926/ [Accessed April 23, 2025].

Goel A, Cantu DA, Guilfoyle J, Chaudhari GR, Newadkar A, Todisco B, de Alba D, Kourdougli N, Schmitt LM, Pedapati E, Erickson CA, Portera-Cailliau C (2018) Impaired perceptual learning in a mouse model of Fragile X syndrome is mediated by parvalbumin neuron dysfunction and is reversible. Nat Neurosci 21:1404–1411 Available at: 10.1038/s41593-018-0231-0.

Goldstone RL (1994) Influences of categorization on perceptual discrimination. J Exp Psychol Gen 123:178–200 Available at: https://pubmed.ncbi.nlm.nih.gov/8014612/ [Accessed May 23, 2025].

Goris J, Braem S, Van Herck S, Simoens J, Deschrijver E, Wiersema JR, Paton B, Brass M, Todd J (2022) Reduced Primacy Bias in Autism during Early Sensory Processing. J Neurosci 42:3989–3999 Available at: https://www.jneurosci.org/content/42/19/3989 [Accessed December 11, 2022].

Goris J, Silvetti M, Verguts T, Wiersema JR, Brass M, Braem S (2021) Autistic traits are related to worse performance in a volatile reward learning task despite adaptive learning rates. Autism 25:440–451 Available at: https://pubmed.ncbi.nlm.nih.gov/33030041/ [Accessed April 25, 2025].

Green SA, Hernandez LM, Bowman HC, Bookheimer SY, Dapretto M (2018) Sensory over-responsivity and social cognition in ASD: Effects of aversive sensory stimuli and attentional modulation on neural responses to social cues. Dev Cogn Neurosci 29:127–139.

Grubb MA, Behrmann M, Egan R, Minshew NJ, Carrasco M, Heeger DJ (2013) Endogenous spatial attention: evidence for intact functioning in adults with autism. Autism Res 6:108 Available at: https://pmc.ncbi.nlm.nih.gov/articles/PMC3661738/ [Accessed April 30, 2025].

Hachen I, Reinartz S, Brasselet R, Stroligo A, Diamond ME (2021) Dynamics of history-dependent perceptual judgment. Nat Commun 2021 121 12:6036- Available at: https://www.nature.com/articles/s41467-021-26104-2 [Accessed February 26, 2026].

Haigh SM (2018) Variable sensory perception in autism. Eur J Neurosci 47:602–609 Available at: https://pubmed.ncbi.nlm.nih.gov/28474794/ [Accessed June 23, 2025].

Harris H, Israeli D, Minshew N, Bonneh Y, Heeger DJ, Behrmann M, Sagi D (2015) Perceptual learning in autism: Over-specificity and possible remedies. Nat Neurosci 18:1574–1576 Available at: https://pubmed.ncbi.nlm.nih.gov/26436903/ [Accessed April 25, 2025].

Hautus, M.J., Macmillan, N.A., & Creelman CD (2021) Detection Theory: A User’s Guide, 3rd ed. Routledge. Available at: 10.4324/9781003203636.

He JL, Williams ZJ, Harris A, Powell H, Schaaf R, Tavassoli T, Puts NAJ (2023) A working taxonomy for describing the sensory differences of autism. Mol Autism 2023 141 14:1–17 Available at: https://molecularautism.biomedcentral.com/articles/10.1186/s13229-022-00534-1 [Accessed July 11, 2024].

He JL, Wodka E, Tommerdahl M, Edden RAE, Mikkelsen M, Mostofsky SH, Puts NAJ (2021a) Disorder-specific alterations of tactile sensitivity in neurodevelopmental disorders. Commun Biol 4:1–15 Available at: 10.1038/s42003-020-01592-y.

He JL, Wodka E, Tommerdahl M, Edden RAE, Mikkelsen M, Mostofsky SH, Puts NAJ (2021b) Disorder-specific alterations of tactile sensitivity in neurodevelopmental disorders. Commun Biol 4 Available at: https://pubmed.ncbi.nlm.nih.gov/33483581/ [Accessed February 28, 2025].

Head J, Helton WS (2014) Sustained attention failures are primarily due to sustained cognitive load not task monotony. Acta Psychol (Amst) 153:87–94 Available at: https://www.sciencedirect.com/science/article/abs/pii/S0001691814002133 [Accessed May 6, 2025].

Järvinen-Pasley A, Wallace GL, Ramus F, Happé F, Heaton P (2008) Enhanced perceptual processing of speech in autism. Dev Sci 11:109–121 Available at: https://pubmed.ncbi.nlm.nih.gov/18171373/ [Accessed June 25, 2025].

Kerzel D, Schönhammer J (2013) Salient stimuli capture attention and action. Atten Percept Psychophys 75:1633–1643 Available at: https://pubmed.ncbi.nlm.nih.gov/23918550/ [Accessed January 26, 2025].

Khachadourian V, Mahjani B, Sandin S, Kolevzon A, Buxbaum JD, Reichenberg A, Janecka M (2023) Comorbidities in autism spectrum disorder and their etiologies. Transl Psychiatry 13:1–7 Available at: https://www.nature.com/articles/s41398-023-02374-w [Accessed April 25, 2025].

Klinger LG, Dawson G (2001) Prototype formation in autism. Dev Psychopathol 13:111–124 Available at: https://pubmed.ncbi.nlm.nih.gov/11346046/ [Accessed April 26, 2025].

Lavie N, Dalton P (2014) Load Theory of Attention and Cognitive Control. Oxford Handb Atten:56–75 Available at: https://academic.oup.com/edited-volume/41256/chapter/350821210 [Accessed April 30, 2025].

Licznerski P et al. (2020) ATP Synthase c-Subunit Leak Causes Aberrant Cellular Metabolism in Fragile X Syndrome. Cell 182:1170–1185.e9 Available at: https://pubmed-ncbi-nlm-nih-gov.proxy.insermbiblio.inist.fr/32795412/ [Accessed July 7, 2024].

Mattioni L, Barbieri A, Grigoli A, Balasco L, Bozzi Y, Provenzano G (2024) Alterations of Perineuronal Net Expression and Abnormal Social Behavior and Whisker-dependent Texture Discrimination in Mice Lacking the Autism Candidate Gene Engrailed 2. Neuroscience 546:63–74 Available at: https://pubmed.ncbi.nlm.nih.gov/38537894/ [Accessed May 19, 2025].

Mercado E, Chow K, Church BA, Lopata C (2020) Perceptual category learning in autism spectrum disorder: Truth and consequences. Neurosci Biobehav Rev 118:689–703 Available at: https://www.sciencedirect.com/science/article/abs/pii/S0149763420305558 [Accessed April 25, 2025].

Micher N, Mazenko D, Lamy D (2024) Unconscious Processing Contaminates Objective Measures of Conscious Perception : Evidence From the Liminal Prime Paradigm. J Cogn:1–16.

Mientjes EJ, Nieuwenhuizen I, Kirkpatrick L, Zu T, Hoogeveen-Westerveld M, Severijnen L, Rifé M, Willemsen R, Nelson DL, Oostra BA (2006) The generation of a conditional Fmr1 knock out mouse model to study Fmrp function in vivo. Neurobiol Dis 21:549–555 Available at: https://pubmed.ncbi.nlm.nih.gov/16257225/ [Accessed June 25, 2025].

Mikkelsen M, Wodka EL, Mostofsky SH, Puts NAJ (2018) Autism spectrum disorder in the scope of tactile processing. Dev Cogn Neurosci 29:140–150 Available at: 10.1016/j.dcn.2016.12.005.

Minshew NJ, Goldstein G, Siegel DJ (1997) Neuropsychologic functioning in autism: Profile of a complex information processing disorder. J Int Neuropsychol Soc 3:303–316 Available at: https://www.cambridge.org/core/journals/journal-of-the-international-neuropsychological-society/article/abs/neuropsychologic-functioning-in-autism-profile-of-a-complex-information-processing-disorder/581818F7F1A05B9413B7BA35EACE2543 [Accessed June 25, 2025].

Minshew NJ, Hobson JA (2008) Sensory sensitivities and performance on sensory perceptual tasks in high-functioning individuals with autism. J Autism Dev Disord 38:1485 Available at: https://pmc.ncbi.nlm.nih.gov/articles/PMC3077539/ [Accessed June 25, 2025].

Mol W, Post S, Lee M, Goel A (2024) Learning Impairments in Fmr1 -/- mice on an Audio-Visual Temporal Pattern Discrimination Task. bioRxiv:2024.09.25.615092 Available at: https://www.biorxiv.org/content/10.1101/2024.09.25.615092v1 [Accessed January 22, 2025].

Molesworth CJ, Bowler DM, Hampton JA (2005) The prototype effect in recognition memory: Intact in autism? J Child Psychol Psychiatry Allied Discip 46:661–672 Available at: https://pubmed.ncbi.nlm.nih.gov/15877770/ [Accessed May 22, 2025].

Mottron L, Dawson M, Soulières I, Hubert B, Burack J (2006) Enhanced perceptual functioning in autism: An update, and eight principles of autistic perception. J Autism Dev Disord 36:27–43.

Murphy G, Groeger JA, Greene CM (2016) Twenty years of load theory—Where are we now, and where should we go next? Psychon Bull Rev 2016 235 23:1316–1340 Available at: https://link.springer.com/article/10.3758/s13423-015-0982-5 [Accessed May 6, 2025].

O’Riordan M, Passetti F (2006) Discrimination in autism within different sensory modalities. J Autism Dev Disord 36:665–675 Available at: https://pubmed.ncbi.nlm.nih.gov/16639532/ [Accessed December 3, 2022].

O’Riordan M, Plaisted K (2001) Enhanced discrimination in autism. Q J Exp Psychol Sect A 54:961–979 Available at: /doi/pdf/10.1080/713756000?download=true [Accessed April 24, 2025].

O’Riordan MA, Plaisted KC, Driver J, Baron-Cohen S (2001) Superior visual search in autism. J Exp Psychol Hum Percept Perform 27:719–730 Available at: https://pubmed.ncbi.nlm.nih.gov/11424657/ [Accessed April 24, 2025].

Orefice LLL, Zimmerman ALL, Chirila AMM, Sleboda SJJ, Head JPP, Ginty DDD (2016) Peripheral Mechanosensory Neuron Dysfunction Underlies Tactile and Behavioral Deficits in Mouse Models of ASDs. Cell 166:299–313 Available at: 10.1016/j.cell.2016.05.033.

Plaisted K, O’riordan M, Baron-Cohen S (1998a) Enhanced Discrimination of Novel, Highly Similar Stimuli by Adults with Autism During a Perceptual Learning Task.

Plaisted K, O’Riordan M, Baron-Cohen S (1998b) Enhanced Visual Search for a Conjunctive Target in Autism: A Research Note. J Child Psychol Psychiatry 39:777–783.

Plaisted KC (2001) Reduced Generalization in Autism: An Alternative to Weak Central Coherence.

Ploog BO (2010) Stimulus overselectivity four decades later: A review of the literature and its implications for current research in autism spectrum disorder. J Autism Dev Disord 40:1332–1349 Available at: https://link.springer.com/article/10.1007/s10803-010-0990-2 [Accessed April 26, 2025].

Puts NAJ, Wodka EL, Tommerdahl M, Mostofsky SH, Edden RAE (2014) Impaired tactile processing in children with autism spectrum disorder. J Neurophysiol 111:1803–1811 Available at: https://pubmed.ncbi.nlm.nih.gov/24523518/ [Accessed December 3, 2022].

Ridderinkhof A, de Bruin EI, van den Driesschen S, Bögels SM (2018) Attention in Children With Autism Spectrum Disorder and the Effects of a Mindfulness-Based Program. J Atten Disord 24:681 Available at: https://pmc.ncbi.nlm.nih.gov/articles/PMC7003152/ [Accessed May 6, 2025].

Robertson CE, Baron-Cohen S (2017) Sensory perception in autism. Nat Rev Neurosci 18:671–684.

Rong Y, Yang CJ, Jin Y, Wang Y (2021a) Prevalence of attention-deficit/hyperactivity disorder in individuals with autism spectrum disorder: A meta-analysis. Res Autism Spectr Disord 83:101759.

Rong Y, Yang CJ, Jin Y, Wang Y (2021b) Prevalence of attention-deficit/hyperactivity disorder in individuals with autism spectrum disorder: A meta-analysis. Res Autism Spectr Disord 83:101759 Available at: https://www.sciencedirect.com/science/article/pii/S1750946721000349 [Accessed May 6, 2025].

Rotschafer SE (2021) Auditory Discrimination in Autism Spectrum Disorder. Front Neurosci 15:1–13.

Samson F, Mottron L, Jemel B, Belin P, Ciocca V (2006) Can spectro-temporal complexity explain the autistic pattern of performance on auditory tasks? J Autism Dev Disord 36:65–76 Available at: https://pubmed.ncbi.nlm.nih.gov/16382329/ [Accessed April 24, 2025].

Sapey-Triomphe LA, Temmerman J, Puts NAJ, Wagemans J (2021a) Prediction learning in adults with autism and its molecular correlates. Mol Autism 12 Available at: https://pubmed-ncbi-nlm-nih-gov.proxy.insermbiblio.inist.fr/34615532/ [Accessed October 12, 2023].

Sapey-Triomphe LA, Timmermans L, Wagemans J (2021b) Priors Bias Perceptual Decisions in Autism, But Are Less Flexibly Adjusted to the Context. Autism Res 14:1134–1146 Available at: https://pubmed-ncbi-nlm-nih-gov.proxy.insermbiblio.inist.fr/33283970/ [Accessed October 12, 2023].

Semelidou O, Gauvrit T, Vandromme C, Cornier A, Saint-Jean A, Le Feuvre Y, Ginger M, Frick A (2024) Stimulus encoding shapes tactile perception and underlies alterations in autism. bioRxiv:2024.08.08.607129 Available at: http://biorxiv.org/content/early/2024/10/08/2024.08.08.607129.abstract.

Shaw KA et al. (2025) Prevalence and Early Identification of Autism Spectrum Disorder Among Children Aged 4 and 8 Years — Autism and Developmental Disabilities Monitoring Network, 16 Sites, United States, 2022. MMWR Surveill Summ 74:1–22 Available at: https://www.cdc.gov/mmwr/volumes/74/ss/ss7402a1.htm [Accessed May 9, 2025].

Shenouda J, Barrett E, Davidow AL, Sidwell K, Lescott C, Halperin W, Silenzio VMB, Zahorodny W (2023) Prevalence and Disparities in the Detection of Autism Without Intellectual Disability. Pediatrics 151 Available at: https://pubmed.ncbi.nlm.nih.gov/36700335/ [Accessed June 25, 2025].

Smallwood J (2013) Penetrating the fog of the decoupled mind: The effects of visual salience in the sustained attention to response task. Can J Exp Psychol 67:32–40.

Soulières I, Mottron L, Giguère G, Larochelle S (2011) Category induction in autism: Slower, perhaps different, but certainly possible. Q J Exp Psychol 64:311–327 Available at: https://pubmed.ncbi.nlm.nih.gov/20623440/ [Accessed April 26, 2025].

Soulières I, Mottron L, Saumier D, Larochelle S (2007) Atypical categorical perception in autism: Autonomy of discrimination? J Autism Dev Disord 37:481–490.

Thye MD, Bednarz HM, Herringshaw AJ, Sartin EB, Kana RK (2018) The impact of atypical sensory processing on social impairments in autism spectrum disorder. Dev Cogn Neurosci 29:151–167.

Venker CE, Mathée J, Neumann D, Edwards J, Saffran J, Ellis Weismer S (2021) Competing Perceptual Salience in a Visual Word Recognition Task Differentially Affects Children With and Without Autism Spectrum Disorder. Autism Res 14:1147–1162 Available at: /doi/pdf/10.1002/aur.2457 [Accessed May 6, 2025].

Zaidel A, Goin-Kochel RP, Angelaki DE (2015) Self-motion perception in autism is compromised by visual noise but integrated optimally across multiple senses. Proc Natl Acad Sci U S A 112:6461–6466 Available at: https://www.pnas.org/doi/abs/10.1073/pnas.1506582112 [Accessed January 5, 2024].

Zetler NK, Cermak SA, Engel-Yeger B, Gal E (2019) Somatosensory discrimination in people with autism spectrum disorder: A Scoping review. Am J Occup Ther 73:1–14.

Zhai J, Li X, Zhou Y, Fan L, Xia W, Wang X, Li Y, Hou M, Wang J, Wu L (2023) Correlation and predictive ability of sensory characteristics and social interaction in children with autism spectrum disorder. Front Psychiatry 14:1056051.

